# Hypermigration of macrophages through the concerted action of GRA effectors on NF-κB/p38 signaling and host chromatin accessibility potentiates *Toxoplasma* dissemination

**DOI:** 10.1101/2024.02.06.579146

**Authors:** Arne L. ten Hoeve, Matias E. Rodriguez, Martin Säflund, Valentine Michel, Lucas Magimel, Albert Ripoll, Tianxiong Yu, Mohamed-Ali Hakimi, Jeroen P. J. Saeij, Deniz M. Ozata, Antonio Barragan

**Affiliations:** Department of Molecular Biosciences, The Wenner-Gren Institute, Stockholm University, Stockholm, Sweden; Program in Bioinformatics and Integrative Biology, University of Massachusetts Medical School, Worcester, MA, USA; Institute for Advanced Biosciences, INSERM U1209, CNRS UMR5309, Université Grenoble Alpes, Grenoble, France; Department of Pathology, Microbiology, and Immunology, University of California Davis, Davis, CA 95616 California, USA

**Keywords:** mononuclear phagocyte, intracellular parasitism, host-pathogen, cell signaling pathway, immune cell migration

## Abstract

Mononuclear phagocytes facilitate the dissemination of the obligate intracellular parasite *Toxoplasma gondii*. Here, we report how a set of secreted parasite effector proteins from dense granule organelles (GRA) orchestrates dendritic cell-like chemotactic and pro-inflammatory activation of parasitized macrophages. These effects enabled efficient dissemination of the type II *T. gondii* lineage, a highly prevalent genotype in humans. We identify novel functions for effectors GRA15 and GRA24 in promoting CCR7-mediated macrophage chemotaxis by acting on NF-κB and p38 MAPK signaling pathways, respectively, with contributions of GRA16/18 and counter-regulation by effector TEEGR. Further, GRA28 boosted chromatin accessibility and GRA15/24/NF-κB-dependent transcription at the *Ccr7* gene locus in primary macrophages. *In vivo*, adoptively transferred macrophages infected with wild-type *T. gondii* outcompeted macrophages infected with a GRA15/24 double mutant in migrating to secondary organs in mice. The data show that *T. gondii*, rather than being passively shuttled, actively promotes its dissemination by inducing a finely regulated pro-migratory state in parasitized human and murine phagocytes via co-operating polymorphic GRA effectors.

**Importance:** Intracellular pathogens can hijack cellular functions of infected host cells to their advantage, for example, for intracellular survival and for dissemination. However, how microbes orchestrate the hijacking of complex cellular processes, such as host cell migration, remains poorly understood. As such, the common parasite *Toxoplasma gondii* actively invades immune cells of humans and other vertebrates and modifies their migratory properties. Here, we show that the concerted action of a number of secreted effector proteins from the parasite, principally GRA15 and GRA24, act on host cell signaling pathways to activate chemotaxis. Further, the protein effector GRA28 selectively acted on chromatin accessibility in the host cell nucleus to selectively boost host gene expression. The joint activities of effectors culminated in pro-migratory signaling within the infected phagocyte. We provide a molecular framework delineating how *T. gondii* can orchestrate a complex biological phenotype, such as the migratory activation of phagocytes to boost dissemination.

## Introduction

Macrophages originate from embryonic progenitors or monocytes and are crucial cells for the innate immune response (1). Being typically sessile cells residing in peripheral tissues, macrophages maintain tissue homeostasis and combat infections with versatile responses, including phagocytosis (2). Conversely, many pathogens have developed strategies to survive and thrive within macrophages, and also manipulate the diverse functions of phagocytes to their advantage (3, 4). Macrophages and dendritic cells (DCs) can be distinguished by transcriptional signatures, for example ZBTB46, IRF4 and BATF3 expression, which also reflects ontology and tissue localization (5).

*Toxoplasma gondii* is an obligate intracellular pathogen commonly carried by humans and many other warm-blooded vertebrates (6). Following oral primary infection, *T. gondii* disseminates broadly in the organism to reach peripheral organs, including the central nervous system. While chronic carriage is chiefly asymptomatic, acute or reactivated infection can cause life-threatening disease in the developing fetus and in immunocompromised individuals (7, 8).

Colonization of the host is mediated by the tachyzoite stage of *T. gondii*. Being obligate intracellular, tachyzoites actively invade host cells (9) in peripheral tissues, including macrophages. The parasite makes use of infected phagocytes for systemic dissemination by a *Trojan horse* mechanism, in a parasite genotype-related fashion (10-12). Mononuclear phagocytes, including principally macrophages, DCs, monocytes and microglia, are induced to migrate via activation of GABAergic signaling and MAP kinase activation (13-16). This migratory activation, termed hypermigratory phenotype (17), implicates secreted parasite effectors (18-21) and impacts motility and chemotaxis of infected phagocytes (22-24).

The invasion of host cells by tachyzoites comprises a discharge of secretory organelles (25, 26). Within parasitized cells, the MYR1 secretory machinery ensures transport of many dense granule proteins (GRAs) across the parasitophorous vacuole (PV), whereafter GRAs can traffic to the host cell cytosol and nucleus (27). GRA proteins present little or no homology to each other and are polymorphic among the clonal lineages of *T. gondii* (type I, II, III) that predominate in Europe and North America. Type II strains are most prevalent in humans and animals used for meat consumption (28, 29).

GRA proteins bear important functions in the immunomodulatory impact of *T. gondii* on the host, such as, activation of the NF-κB pathway and MAP kinase signaling in macrophages (30, 31) and chromatin remodeling, which impact transcription in the host cell (32, 33). Notably, recent findings identified a role for the chromatin remodeler-interacting effector GRA28 (type I) on the migratory activation of phagocytes. Among the features of this activation lay the expression of chemokine receptor CCR7, onset of chemotaxis and systemic migration by parasitized macrophages in mice (21).

Here, we investigated the molecular machinery that imparts a DC-like migratory activation on parasitized macrophages. We show that the concerted action of polymorphic effectors regulates the pro-migratory activation of parasitized macrophages. Specifically, a set of secreted GRA proteins impacts NF-κB/p38 MAPK signaling and host chromatin accessibility to co-operatively promote CCR7-driven chemotaxis in infected macrophages and, thereby, potentiate parasite dissemination.

## Results

### The effector GRA15 mediates a DC-like migratory activation in *T. gondii*-infected macrophages

The infection of macrophages by *T. gondii* tachyzoites (type I) initiates a migratory activation, which encompasses expression of DC-associated transcription factors, phenotypical changes and onset of CCR7-dependent chemotaxis (21). Because NF-κB positively regulates CCR7 expression in DCs (34) and the *T. gondii* protein GRA15 (type II) activates NF-κB (30), we hypothesized that GRA15 contributes to the migratory activation of parasitized macrophages. Bone marrow-derived macrophages (BMDMs) challenged with type II *T. gondii* tachyzoites (wild-type PRU) (**Figure 1A**) dramatically upregulated the expression of *Ccr7* mRNA (**Figure 1B**). Interestingly, BMDMs challenged with GRA15-deficient tachyzoites (PRUΔ*gra15*) consistently exhibited a significantly decreased *Ccr7* expression (**Figure 1B**), compared with wild-type-challenged BMDMs at similar infection frequencies (**Fig S1A**). To functionally assess the putative impact of the induced *Ccr7*, we performed chemotaxis assays with the CCR7-ligand chemokine CCL19. Notably, wild-type PRU-infected BMDMs, but not bystander BMDMs, displayed a distinct migratory response towards the CCL19 source (**Figure 1C**). In sharp contrast, chemotaxis to CCL19 was dramatically reduced in BMDMs challenged with PRUΔ*gra15* (**Figure 1C)**, while random hypermotility (22) with elevated velocity was maintained (**Figure S1B**). Collectively, the findings suggest a role for GRA15 in mediating CCR7-dependent chemotaxis, without measurable effects on bystander cells or on hypermotility (13).

**Figure 1.**
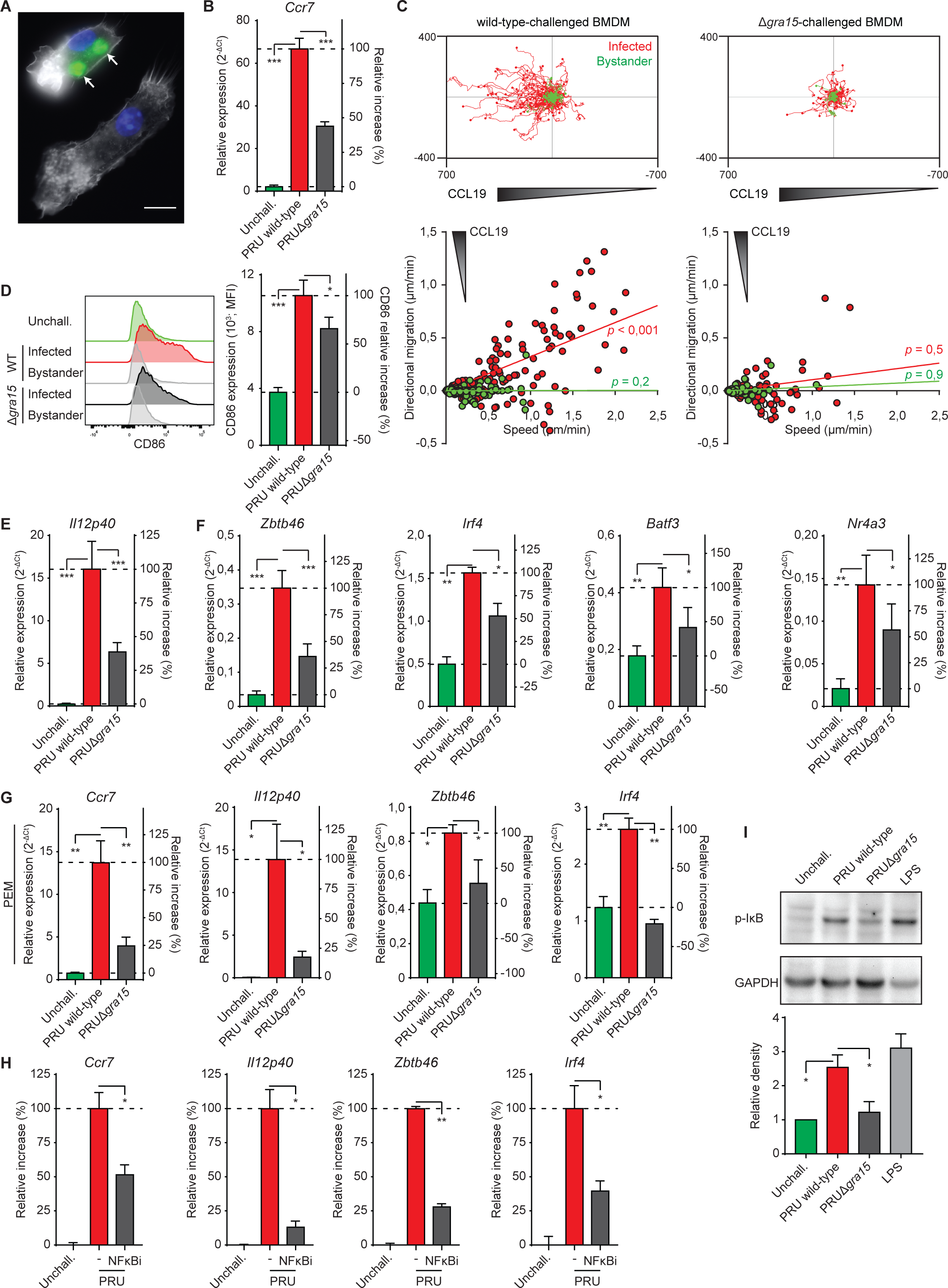
Phenotypes *T. gondii*-infected macrophages upon GRA15-deficiency and NF-κB inhibition. (**A**) Representative micrograph shows primary bone marrow-derived macrophages (BMDMs) stained for F-actin (phalloidin Alexa Fluor 594, white) and nuclei (DAPI, blue). Arrows indicate two intracellular vacuoles with replicating GFP-expressing type II *T. gondii* tachyzoites (green) 18 h post-challenge. Lower by-stander cell is uninfected. Scale bar = 10 µm. (**B**) qPCR analysis of *Ccr7* cDNA from BMDMs challenged for 18 h with freshly egressed *T. gondii* type II wild-type and GRA15-deficient (Δ*gra15*) tachyzoites (PRU; MOI 2). For reference, macrophages were incubated in complete medium, unchallenged (unchall.). Displayed are relative expression (2^-ΔCt^) and the increase in expression relative to wild-type (100%) and unchallenged (0%) conditions (mean + SEM; n=4 independent experiments). (**C**) Motility plots depict the displacement of BMDMs challenged with freshly egressed *T. gondii* type II wild-type and GRA15-deficient (Δ*gra15*) tachyzoites (PRU; MOI 1) over 14 h in a collagen matrix with a CCL19 gradient as detailed in Methods (scale indicates µm; n=3). For each condition, directional migration (µm/min) towards the CCL19 source and speed (µm/min) of individual cells are displayed in graphs, with linear regression lines. Infected cells (GFP^+^, red) and non-infected bystander cells (GFP^-^, green) were analyzed. For each condition, p-values indicate the directional migration compared to hypothetical zero directionality (one-sample permutation test). (**D**) Flow cytometric analysis of anti-CD86 staining on BMDMs challenged for 18 h with freshly egressed GFP-expressing *T. gondii* type II wild-type (WT) and GRA15-deficient (Δ*gra15*) tachyzoites (PRU; MOI 1) or left unchallenged. Infected (GFP^+^) and bystander cells (GFP^-^) were analyzed. Bar graph displays the MFI and the increase in expression relative to wild-type (100%) and unchallenged (0%) conditions (mean + SEM; n=5). (**E**) and (**F**) qPCR analyses of *Il12p40* (E) or *Zbtb46*, *Irf4* and *Nr4a3* (F) cDNA from BMDMs challenged and displayed as in (B), n=4. (**G**) qPCR analyses of *Ccr7*, *Il12p40*, *Zbtb46* and *Irf4* cDNA from resident peritoneal macrophages (PEMs) challenged and displayed as in (B), n=3. (**H**) qPCR analyses of *Ccr7*, *Il12p40*, *Zbtb46* and *Irf4* cDNA from BMDMs challenged for 18h with GFP-expressing *T. gondii* type II wild-type tachyzoites with or without JSH-23 treatment (NFκBi). Displayed is the increase in expression relative to untreated unchallenged (0%) and wild-type (100%) challenged conditions (mean + SEM; n=3). (**I**) Western blot analysis of phospho-IκBα ser32/36 (p-IκB) levels in BMDMs challenged for 5 h with freshly egressed *T. gondii* type II wild-type or GRA15-deficient (Δ*gra15*) tachyzoites (PRU, MOI 3) or LPS (10 ng/mL) or left unchallenged (unchall.). Bar graph displays the relative density of specific p-IκB signal relative to specific GAPDH signal (mean + SEM; n=3). Statistical comparisons were made with ANOVA and Dunnett’s post-hoc (B, D-I) or one-sample permutation tests (B; * p ≤ 0,05, ** p ≤ 0,01, *** p ≤ 0,001, ns p > 0,05).

Next, we assessed the impact of GRA15 on the expression of co-stimulatory molecules (CD40/80/86), MHCII and *Il12p40* because they are also regulated by NF-κB (35, 36). *T. gondii* wild-type (PRU)-infected BMDMs, but not bystander BMDMs, robustly upregulated CD86 expression (**Figure 1D**), while CD40, CD80 and MHCII were minorly affected (**Figure S1D** and **S1E**). Notably, PRUΔ*gra15*-infected BMDMs expressed reduced CD80 and CD86 levels compared with wild-type-infected BMDMs. Further, wild-type-challenged BMDMs upregulated *Il12p40* expression, which was significantly reduced in PRU*Δgra15*-challenged BMDMs (**Figure 1E**). Finally, we determined the contribution of GRA15 to the expression of DC-associated transcription factors, normally not expressed or expressed at low level in macrophages (21). Consistently, BMDMs challenged with wild-type tachyzoites elevated the expression of *Zbtb46*, *Irf4*, *Batf3* and *Nr4a3* and GRA15-deficiency significantly reduced the relative expression of these genes (**Figure 1F**). Data were confirmed using primary peritoneal macrophages (**Figure 1G**). In contrast, early growth response 1 (*Egr1*) expression, known to be GRA24-dependent (31, 37), was significantly enhanced by GRA15-deficiency (**Figure S1F**). Next, cells were treated with separate NF-κB inhibitors targeting nuclear translocation of NF-κB or activator IκB kinase (IKK) (38, 39). Similar for both inhibitors, treated *T. gondii*-challenged BMDMs expressed significantly lower levels of *Ccr7, Il12p40, Zbtb46* and *Irf4* than their untreated counterparts (**Figure 1H** and **S1G**), thus mimicking the effects of GRA15-deficiency. Finally, we confirmed a GRA15-dependent S32/36 phosphorylation of the NF-κB cytoplasmic anchor IκB in *T. gondii*-challenged BMDMs (**Figure 1I**). We conclude that in *T. gondii* type II (PRU)-infected macrophages, GRA15 contributes to the inductions of CCR7-dependent chemotaxis, *Il12p40* and DC-associated transcription factors in an NF-κB-dependent manner.

### Impact of the MYR1-dependent effector GRA24 on the migratory activation of macrophages

Several *T. gondii*-derived proteins that modulate host cell signaling rely on the MYR1 secretory pathway for translocation to the host cell cytosol and nucleus (40). We recently identified a role for the MYR1-translocated effector GRA28 in the induction of DC-like migratory activation of macrophages by the type I RH strain (21). However, the facts that type I RH strain lacks a functional GRA15 (30) and that the reduction of macrophage activation upon GRA15-deficiency (type II) was partial (Figure 1), motivated investigation of additional putative effectors. First, we confirmed that the inductions of *Ccr7*, *Il12p40*, *Zbtb46*, *Nr4a3, Irf4* and *Batf3* were significantly reduced in macrophages challenged with type II MYR1-deficient *T. gondii* (PRUΔ*myr1*) (**Figure 2A**). Because p38 mitogen-activated protein kinase (MAPK) can act as a positive regulator of CCR7 expression in DCs (41), we hypothesized a contribution of the MYR1-secreted p38-activating effector GRA24 (27, 31) to the induction of *Ccr7* and CCR7-dependent chemotaxis in infected macrophages. In line with this idea, GRA24-deficient (PRUΔ*gra24*) tachyzoites induced significantly lower levels of *Ccr7* (∼50% reduction) compared with wild-type *T. gondii* and reconstitution (PRUΔgra24+GRA24) resulted in elevated *Ccr7* expression (**Figure 2B**). Consistent with the transcriptional data, PRUΔ*gra24*-infected BMDMs did not display chemotaxis towards CCL19, while chemotaxis was recovered in GRA24-reconstituted PRUΔ*gra24 T. gondii* (**Figure 2C**). Similar to *Ccr7*, the expressions of *Il12p40*, *Zbtb46*, *Irf4*, *Batf3* and *Nr4a3* were reduced upon challenge with PRUΔ*gra24* (**Figure 2D**). Finally, we confirmed that, in *T. gondii*-challenged BMDMs, cytoplasmic phosphorylated p38 (p-p38) MAPK levels were maintained in a GRA24-dependent manner (**Figure 2E**). Together, the data implicate GRA24/p38 in the migratory activation of macrophages and motivated a further investigation of MAPK signaling.

**Figure 2.**
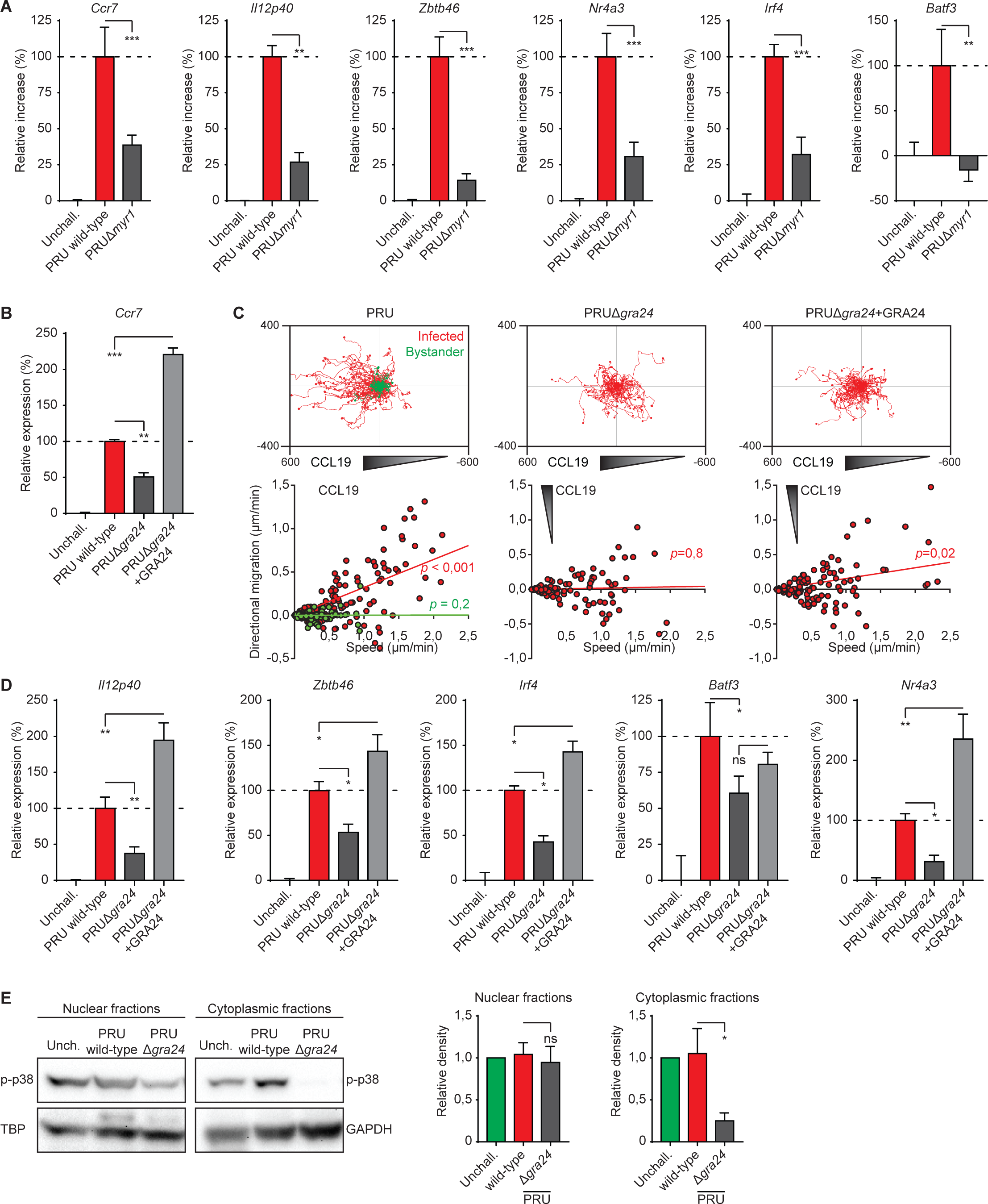
Phenotypes of *T. gondii*-infected macrophages upon MYR1- and GRA24-deficiency. (**A**) qPCR analyses of *Ccr7*, *Il12p40*, *Zbtb46*, *Nr4a3*, *Irf4* and *Batf3* cDNA from BMDMs challenged for 18 h with freshly egressed *T. gondii* type II wild-type and MYR1-deficient (Δ*myr1*) tachyzoites (PRU; MOI 2) or left unchallenged (unchall.). Bar graphs display the increase in expression relative to untreated unchallenged (0%) and wild-type (100%) challenged conditions (mean+SEM; n=4). (**B**) qPCR analysis of *Ccr7* cDNA from BMDMs challenged for 18h with *T. gondii* type II wild-type, GRA24-deficient (Δ*gra24*) or GRA24-reconstituted (Δ*gra24*+GRA24) tachyzoites (PRU; MOI 2) or left unchallenged (unchall.), displayed as in (A), n=3. (**C**) Motility plots depict the displacement of BMDMs challenged with freshly egressed *T. gondii* type II wild-type and GRA24-deficient (Δ*gra24*) tachyzoites (PRU; MOI 1) over 14 h in a collagen matrix with a CCL19 gradient as detailed in Methods (scale indicates µm; n=3). For each condition, directional migration (µm/min) towards the CCL19 source and speed (µm/min) of individual cells are displayed in graphs, with linear regression lines. Infected cells (GFP^+^, red) and non-infected bystander cells (GFP^-^, green) were analyzed. For each condition, p-values indicate the directional migration compared to hypothetical zero directionality (one-sample permutation test). (**D**) qPCR analyses of *Il12p40, Zbtb46, Irf4, Batf3* and *Nr4a3* of BMDMs challenged and displayed as in (B). (**E**) Western blot analysis of p-p38 (Thr180/Tyr182) expression in cytoplasm- and nucleus-enriched fractions of BMDMs challenged for 5 h with wild-type and GRA24-deficient (Δ*gra24*) *T. gondii* type II tachyzoites (PRU, MOI 3). Bar graphs display relative density of p-p38 signal relative to TBP or GAPDH signals (mean+SEM; n=5). Statistical comparisons were made with ANOVA and Dunnett’s post-hoc tests (* p ≤ 0,05, ** p ≤ 0,01, *** p ≤ 0,001, ns p > 0,05).

### Ribosomal S6 kinase (RSK) regulates the migratory activation of *T. gondii*-infected macrophages

Because MAPK signaling is strongly linked to cell motility and chemotaxis, we evaluated the impact of pharmacological inhibition of principal MAPK pathways (ERK1/2, ERK5, p38, JNK) on the DC-like migratory activation of macrophages. Inhibition of p38 MAPK significantly reduced expression of *Ccr7*, *Il12p40*, *Zbtb46* and *Irf4* in PRU-challenged BMDMs (**Figure 3A**), while MEK5 (ERK5) inhibition reduced *Il12p40*, *Zbtb46* and *Irf4* expression (**Figure 3B**). Notably, the transcriptional inductions were conversely amplified by MEK1/2 (ERK1/2) inhibition (**Figure 3A)**, in a GRA24-independent manner (**Figure S2A**), and in line with reported inhibitory effects of ERK1/2 on IL-12 production (42). Yet, inhibition of Ca^2+^/calmodulin-dependent activation of MEK1/2 or ERK1/2-dimerization did not recapitulate this amplification (**Figure S2B**). Consistent with above and Δ*gra24* data (Figure 2C), p38 inhibition abolished chemotaxis, which was maintained upon MEK1/2 inhibition (**Figure S2C**). Jointly, the findings implicate p38 MAPK signaling in the GRA24-driven chemotactic responses of macrophages.

**Figure 3.**
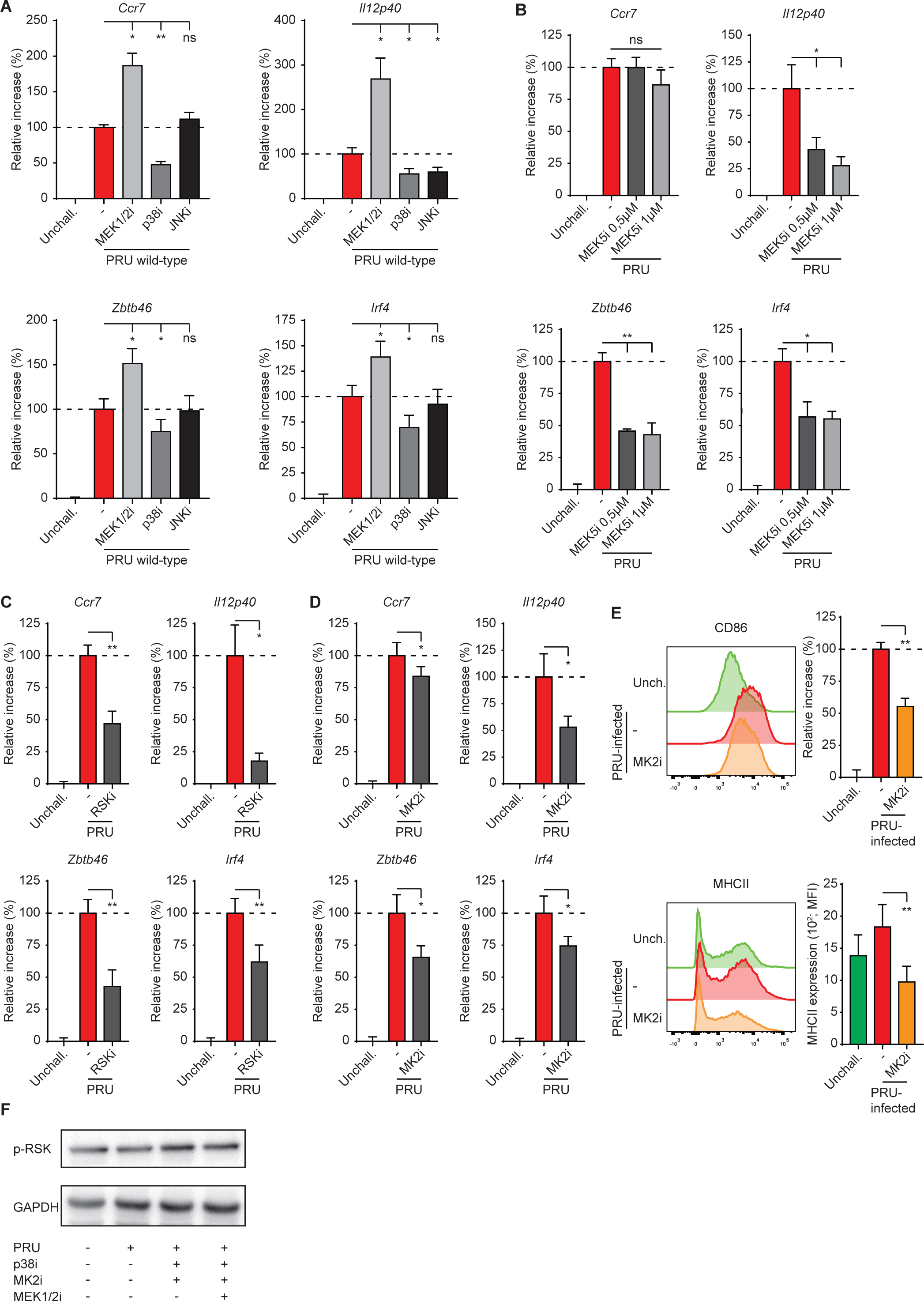
Migratory activation in *T. gondii*-infected macrophages involves MAPK-associated kinases. (**A**) and (**B**) qPCR analyses of *Ccr7*, *Il12p40*, *Zbtb46* and *Irf4* cDNA from BMDMs challenged for 18h with GFP-expressing *T. gondii* type II wild-type tachyzoites with (A) Trametinib (MEK1/2i), BIRB 796 (p38i) or JNK-IN-8 (JNKi) treatments, or (B) BIX02189 (MEK5i) treatment. Displayed is the increase in expression relative to untreated unchallenged (unchall., 0%) and wild-type (100%) challenged conditions (mean + SEM, n=4 (A) and (B)). (**C**) and (**D**) qPCR analyses of *Ccr7*, *Il12p40*, *Zbtb46* and *Irf4* cDNA from BMDMs challenged for 18h with GFP-expressing *T. gondii* type II wild-type tachyzoites with or without BRD7389 (RSKi; C) or MK2-IN-1 (MK2i; D) treatment. Displayed as in (A), n=4 (C) or 3 (D). (**E**) Flow cytometric analysis of anti-CD86 and MHCII staining on BMDMs challenged for 18 h with freshly egressed GFP-expressing *T. gondii* type II wild-type tachyzoites (PRU; MOI 1), with or without MK2-IN-1 (MK2i) treatment, or left unchallenged. Infected (GFP^+^) cells were analyzed for challenged conditions. Bar graph displays the increase in expression, as in (A), n=4. (**F**) Western blot analysis of p-RSK (S380/386) expression in lysates of BMDMs challenged for 5h with wild-type *T. gondii* type II tachyzoites (PRU, MOI 3) in the presence of indicated inhibitors. Representative of 2 experiments. Statistical comparisons were made with ANOVA and Dunnett’s post-hoc tests (A), paired t-test (C-E) and Spearman correlation (B; * p ≤ 0,05, ** p ≤ 0,01, *** p ≤ 0,001, ns p > 0,05).

Next, we tested the impact of MAPK-activated kinases RSK and MK2, two crucial downstream effectors of MAPK signaling (43). Inhibition of RSK nearly abolished the induction of *Il12p40* and significantly reduced expression of *Ccr7*, *Zbtb46* and *Irf4* in wild-type PRU-challenged BMDMs (**Figure 3C**). Inhibition of MK2 also reduced *Il12p40, Ccr7*, *Zbtb46* and *Irf4* expression in *T. gondii*-challenged BMDMs (**Figure 3D**) and reduced MHCII and CD86 expression on *T. gondii*-infected BMDMs (**Figure 3E**). In BMDCs and BMDMs, RSK can be activated by ERK1/2 and, independently, p38-MK2 signaling (44). To confirm that RSK was indeed activated via p38-MK2 or ERK, we evaluated the RSK phosphorylation at Ser380/386. We found that p-RSK was readily detectable in unchallenged BMDMs, which was maintained in cells challenged with wild-type (PRU) or GRA24-deficient tachyzoites (**Figure S2D** and **S2E**). However, p-RSK levels were not notably reduced by combinations of p38/MK2 or p38, MK2 and MEK1/2 inhibition in PRU-challenged BMDMs (**Figure 2F**), making its activation mechanism elusive. Finally, because *Ccr7* expression can also be regulated by AP-1 (Fos/Jun) and ETS PU.1 transcription factors, which in turn are regulated by p38 MAPK (45, 46), we applied pharmacological inhibition. AP-1 and PU.1 inhibition did not impact expression of *Ccr7, Zbtb46* and *Irf4* while expression of *Il12p40* was significantly down- and upregulated, respectively (**Figure S3A** and **S3B**). We conclude that, in *T. gondii*-infected BMDMs, p38-MK2 and RSK contribute to the induction of CCR7-dependent chemotaxis and DC-associated transcription factors, without across-the-board measurable involvement of AP-1 or PU.1.

### Additive effects of GRA15–NF-κB and GRA24–p38 signaling on the migratory activation of macrophages

Because macrophages infected with either GRA15 or GRA24 single knockout parasites had reduced but not abolished *Ccr7* expression (Figures 1 and 2), we hypothesized that collective effects were in play. BMDMs challenged with a GRA15/24 double mutant (Δ*gra15*Δ*gra24*) had significantly reduced *Ccr7* expression compared with Δ*gra15-* and Δ*gra24-*challenged BMDMs (**Figure 4A**). Expression levels of *Il12p40*, *Zbtb46*, *Irf4*, *Nr4a3* and *Batf3* were, however, not significantly further reduced by double-deficiency of GRA15/24 (**Figure 4A** and **4B**). Consistent with its GRA24-dependency, *Egr1* expression was abolished in PRUΔ*gra24* and Δ*gra15*Δ*gra24*-challenged BMDMs while GRA15-deficiency significantly elevated mRNA levels of *Egr1* (**Figure 4C** and **S1F**). Next, we assessed the implications of NF-κB (GRA15) and p38 (GRA24) signaling by combined pharmacological inhibition. Individually, inhibitors partially inhibited *Ccr7*, *Il12p40*, *Zbtb46* and *Irf4* expression in PRU-challenged macrophages with an overall superior effect upon NF-κB inhibition, while combined inhibition of NF-κB and p38, expectedly, mirrored the effects of challenge with Δ*gra15*Δ*gra24* parasites (**Figure 4D**).

**Figure 4.**
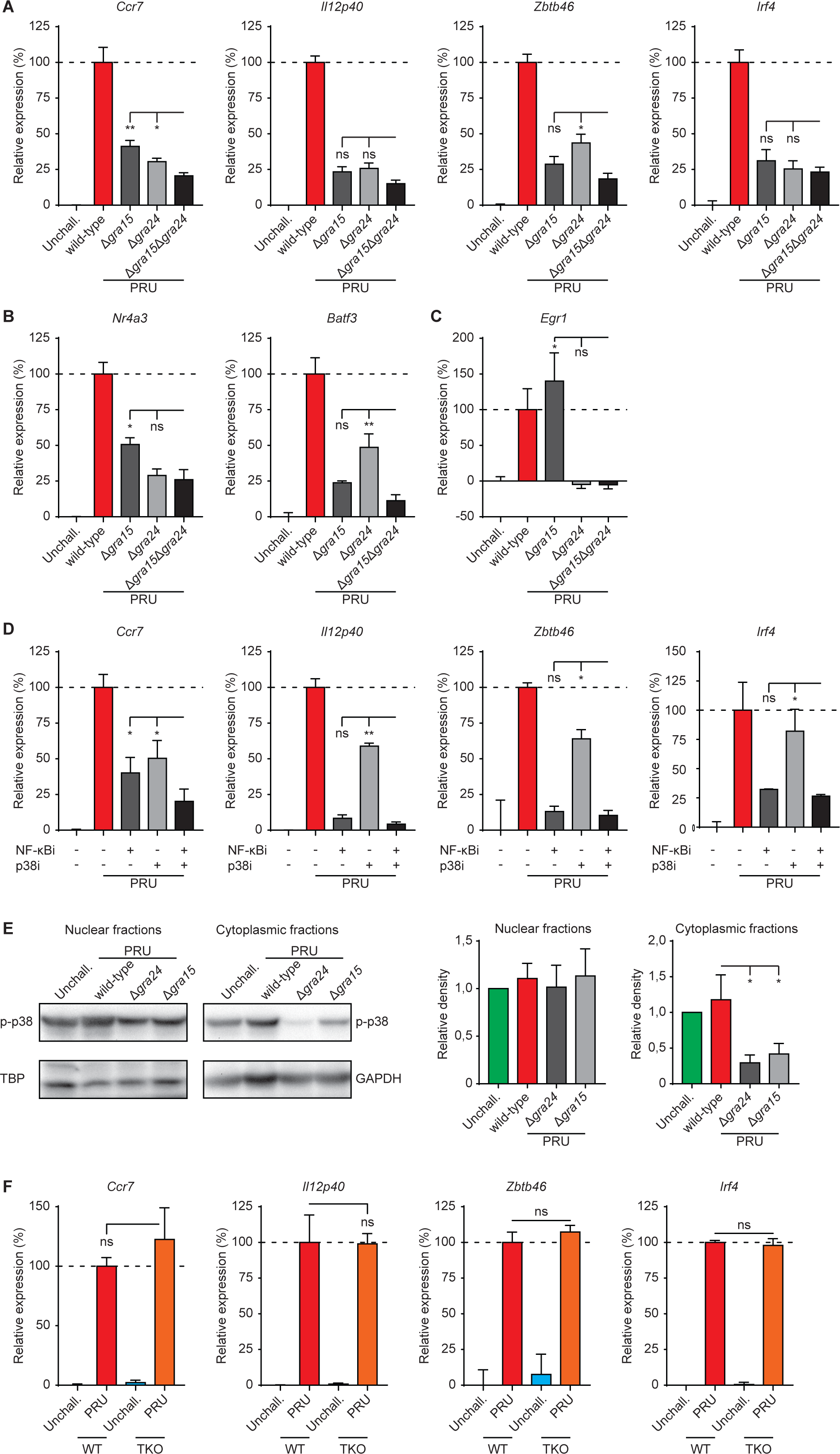
Effects of GRA15/24 double deficiency, NF-κB/p38 MAK inhibition and canonical NF-κB signaling on the activation of *T. gondii*-infected macrophages. (**A**), (**B**) and (**C**) qPCR analyses of (A) *Ccr7*, *Il12p40*, *Zbtb46*, *Irf4*, (B) *Nr4a3*, *Batf3* and (C) *Egr1* cDNA from BMDMs challenged with *T. gondii* type II PRU (wild-type), GRA15 (Δ*gra15*), GRA24 (Δ*gra24*) or GRA15/24 double mutant (Δ*gra15*Δ*gra24*) tachyzoites (18h, MOI 2). Displayed is the increase in expression relative to untreated unchallenged (unchall., 0%) and wild-type (100%) challenged conditions (mean + SEM, n=4). (**D**) qPCR analyses of *Ccr7*, *Il12p40*, *Zbtb46* and *Irf4* cDNA from BMDMs challenged for 18h with *T. gondii* type II wild-type tachyzoites in presence of NFκB inhibitor (JSH-23, NFκBi), p38 inhibitor (BIRB 796, p38i) or combined treatment. Displayed is the increase in expression relative to untreated unchallenged (0%) and wild-type (100%) challenged conditions (mean + SEM; n=3). (**E**) Western blot analysis of p-p38 (Thr180/Tyr182) expression in nucleus- and cytoplasm-enriched fractions of BMDMs challenged with PRU (wild-type), Δ*gra24* or Δ*gra24* tachyzoites (5 h, MOI 3). Bar graphs display relative density of p-p38 signal relative to TBP or GAPDH signals (mean+SEM; n=4). (**F**) qPCR analyses of *Ccr7*, *Il12p40*, *Zbtb46* and *Irf4* cDNA from wild-type BMDMs (WT) or Myd88^-/-^ Ticam^-/-^ Mavs^-/-^ BMDMs (TKO) challenged for 18h with *T. gondii* type II wild-type tachyzoites (MOI 2). Displayed is the increase in expression relative to wild-type (100%) (mean + SEM; n=3) Statistical comparisons were made with ANOVA and Dunnett’s post-hoc tests (* p ≤ 0,05, ** p ≤ 0,01, *** p ≤ 0,001, ns p > 0,05).

GRA15 activates NF-κB through tumor necrosis factor receptor-associated factors (TRAFs) (47). In turn, TRAF6 may also mediate p38 activation in cells (48). Consistent with this idea, Western blotting revealed significantly lowered cytoplasmic p-p38 levels in BMDMs challenged with Δ*gra24* and, interestingly, also with Δ*gra15* parasites (**Figure 4E**), indicating an impact of GRA15 on p-p38. Finally, to determine the contribution of NF-κB activation via pattern recognition receptors (PRRs), we challenged macrophages derived from Myd88^-/-^ Ticam^-/-^ Mavs^-/-^ mice (49). Interestingly, the responses of *T. gondii*-challenged mutant BMDMs approximated the responses by wild-type BMDMs (**Figure 4F**), indicating a minor or non-significant contribution of NF-κB activation via PRRs to this phenotype. In sharp contrast, *Il12p35/p40* responses to LPS were undetectable in Myd88^-/-^ Ticam^-/-^ Mavs^-/-^ BMDMs (**Figure S4**). Jointly, the data show that signaling linked to effectors GRA15 and GRA24 likely co-operates in the migratory activation and the DC-like transcriptional impact on macrophages via NF-κB signaling and p38 MAPK signaling, including a cross-regulation between the two pathways (50, 51). Yet, the intriguing finding that the effects on *Ccr7* were strongly reduced (∼80%), but not strictly abolished, by GRA15/24 double-deficiency or by combined pharmacological inhibition motivated a further exploration of additional putative effectors.

### Counteractive effects of parasite effector TEEGR and contributions of GRA16/18 to the activation of macrophages

Having established that NF-κB signaling contributes to the upregulation of *Ccr7*, *Il12p40* and DC-associated transcription factors with pro-migratory effects, we investigated the effector TEEGR because of its reported counteracting effects on NF-κB-dependent gene induction (52). Contrasting with the effects of GRA15 deficiency, BMDMs challenged with TEEGR-deficient type II tachyzoites (PRUΔ*teegr*) consistently expressed significantly higher levels of *Ccr7*, *Il12p40* and *Zbtb46* compared with wild-type-challenged macrophages (**Figure 5A**). Next, we tested the effector GRA16 in a similar fashion because of its reported indirect inhibitory effect on NF-κB in an epithelial-like cell line (53). Surprisingly, GRA16-deficient type II tachyzoites (PRUΔ*gra16*) reduced the induction of *Ccr7*, *Il12p40* and *Zbtb46* compared with wild-type *T. gondii* (PRU) in macrophages (**Figure 5B**). This may suggest that inhibition of NF-κB signaling is not the main role of GRA16 in the migratory activation of infected macrophages. Finally, because GRA18 stimulates expression of genes related to the inflammatory response and chemotaxis (54), we addressed its putative contribution. Indeed, GRA18-deficient (Δ*gra18*) tachyzoites induced significantly lower expression levels of *Ccr7*, *Il12p40*, *Zbtb46* and *Irf4* compared with wild-type (PRU) parasites in BMDMs (**Figure 5C**). GRA18 interacts with phosphatase complex PP2A-B56 and the negative regulator of GSK3α/β β-catenin (54). Pharmacological GSK inhibition further reduced expression of *Ccr7*, *Zbtb46* and *Irf4* in PRU*Δgra18*-challenged BMDMs. However, in wild-type -challenged macrophages, GSK3 inhibition significantly reduced *Irf4*, but not *Ccr7* or *Zbtb46* expression (**Figure S5**), indicating additional regulation. Together, the data indicate that parasite effector TEEGR counteracts pro-migratory signaling, likely by NF-κB repression. Conversely, GRA16 and GRA18 promote signaling consistent with pro-migratory activation of infected macrophages.

**Figure 5.**
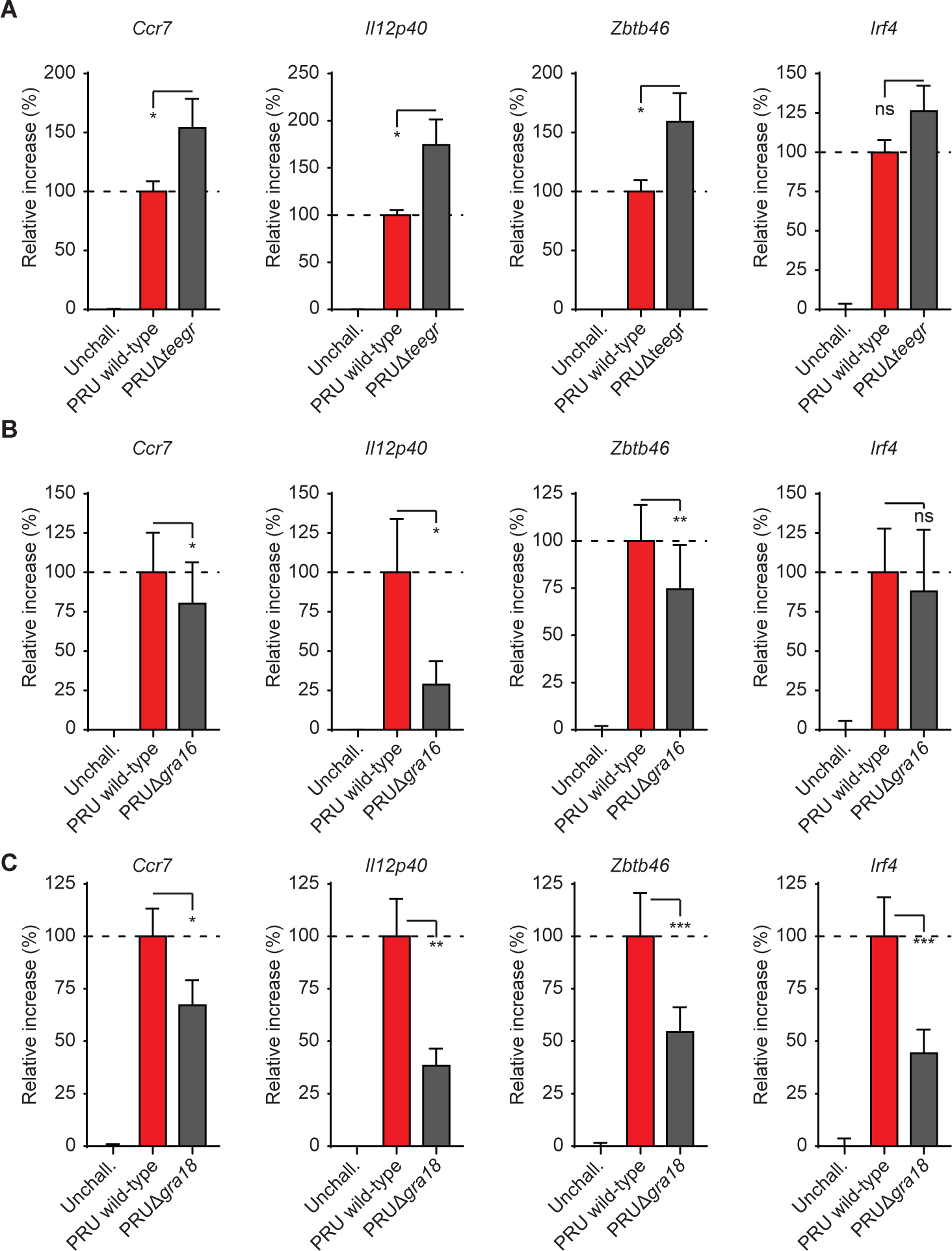
Transcriptional impacts of TEEGR, GRA16 and GRA18 mutants on macrophage activation. (**A**), (**B**) and (**C**) qPCR analyses of *Ccr7*, *Il12p40*, *Zbtb46* and *Irf4* cDNA from BMDMs challenged for 18h (MOI2) with *T. gondii* type II PRU (wild-type), (A) TEEGR-deficient mutant (Δ*teegr*), (B) GRA16-deficient mutant (Δ*gra16*) or (C) GRA18-deficient mutant (Δ*gra18*). Displayed is the increase in expression relative to unchallenged (unchall., 0%) and wild-type (100%) challenged conditions (mean + SEM, n=5 (A; B) and n=7(C).

### Selective impact of GRA28 on chromatin accessibility at the *Ccr7* locus and other gene loci implicated in the activation of macrophage migration

Gene expression is linked to chromatin accessibility, which is modulated by chromatin remodeling complexes, such as SWI/SNF (55). Analyses of publicly available datasets (56) using Assay for Transposase-Accessible Chromatin with sequencing (ATAC-seq) revealed accessible chromatin around the promoters of *Ccr7*, *Il12p40* and *Zbtb46* genes in different types of DCs but not in macrophages (**Figure S6A**). We previously showed that GRA28 (type I) interacts with the SWI/SNF complex and binds to chromatin (21). Together, this motivated an assessment of chromatin accessibility for genes linked to the migratory activation of BMDMs using ATAC-seq, in the presence or absence of GRA28 (type II). Our analysis detected 26,663 open chromatin peaks uniquely found in BMDMs challenged with wild-type (PRU) tachyzoites compared with unchallenged BMDMs (**Figure 6A** and **S6B**). Further, among these 26,663 peaks, 11,067 peaks were not detected in PRUΔ*gra28*-challenged BMDMs, underlying the involvement of GRA28 in chromatin accessibility (**Figure 6A**). Importantly, around the transcription start site of *Ccr7*, we found a notable increase in accessible chromatin in wild-type PRU-infected BMDMs compared with unchallenged BMDMs, which is a typical feature of DCs **(Figure 6B**). Supporting the idea that GRA28 drives chromatin accessibility, we observed a reduction in accessible chromatin in PRUΔ*gra28*-infected BMDMs **(Figure 6B**). In sharp contrast, increases in chromatin accessibility were not detected around promoters of *Ccr2*, *Ccr5* and *Cx3cr1* genes, coding for other chemokine receptors in phagocytes (**Figure S7A**). In line with ATAC-seq data, GRA28-deficiency (PRUΔ*gra28*) nearly abolished the induction of *Ccr7,* determined by RT-qPCR (**Figure 6C**). Finally, chemotaxis to CCL19 was undetectable in BMDMs challenged with PRUΔ*gra28* strain (**Figure 6D**), functionally corroborating the findings.

**Figure 6.**
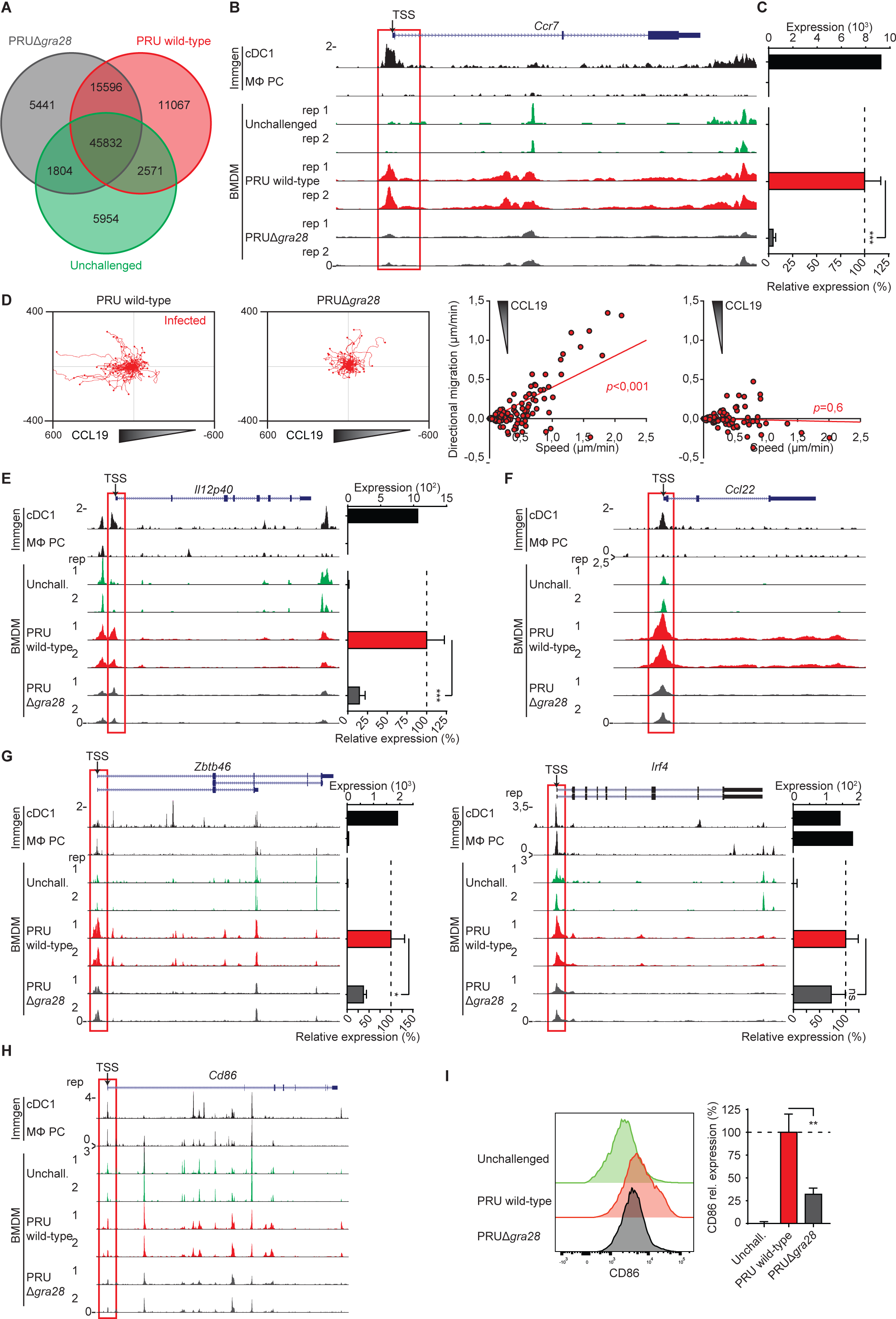
Chromatin accessibility and phenotypes of macrophages challenged with wild-type and GRA28-deficient *T. gondii* tachyzoites. (**A**) Venn diagram shows numbers of identified ATAC-seq peaks in unchallenged BMDMs and BMDMs challenged for 18h with *T. gondii* type II wild-type or GRA28-deficient (Δ*gra28*) tachyzoites (PRU; MOI 2). (**B**) Genome tracks show ATAC-seq peak intensities, normalized to uniquely aligned reads (y-axis), around the promoter of *Ccr7* gene. For BMDMs, ATAC-seq signal is from 2 separate biological replicates per condition. Upper tracks show peak signal from dendritic cells (cDC1) and peritoneal cavity macrophages (M<l PC) extracted from Immgen publicly available dataset. Red outline indicates region of interest near the transcription start site (TSS). (**C**) Upper bar graph shows *Ccr7* expression by cDC1 and M<l PC, quantified by RNA-seq (Immgen publicly available dataset). Lower bar graph shows qPCR analysis of *Ccr7* cDNA from BMDMs challenged for 18h with *T. gondii* type II wild-type or GRA28-deficient (Δ*gra28*) tachyzoites (PRU; MOI 2) or left unchallenged. The increase in expression relative to unchallenged (unchall., 0%) and wild-type (100%) challenged conditions is displayed (mean + SEM, n=5). (**D**) Motility plots depict the displacement of BMDMs challenged with CMTMR-stained *T. gondii* type II wild-type and GRA28-deficient (Δ*gra28*) tachyzoites (PRU; MOI 1) in a CCL19 gradient, as detailed in Methods (scale indicates µm). For each condition, directional migration (µm/min) towards the CCL19 source and speed (µm/min) of individual cells are displayed in graphs, with linear regression lines. Infected cells (CMTMR^+^) were analyzed from 3 independent experiments. For each condition, p-values indicate the directional migration compared to hypothetical zero directionality (one-sample permutation test). (**E-I**) ATAC-seq genome tracks for (E) *Il12p40,* (F) *Ccl22,* (G) *Zbtb46,* (H) *Irf4* and (I) *CD86* in cDC1 and MΦ PC as in (B), and transcriptional analyses as in (C). (**J**) Flow cytometric analysis of anti-CD86 staining on BMDMs challenged for 18 h with CFSE-stained *T. gondii* type II wild-type tachyzoites (PRU; MOI 1). Infected (CFSE^+^) cells were analyzed for challenged conditions. Bar graph displays expression related to wild-type (mean + SEM, n=4). Statistical comparisons were made with ANOVA and Dunnett’s post-hoc tests (C, E, G, H, J, * p ≤ 0,05, ** p ≤ 0,01, ** p ≤ 0,001, ns p > 0,05).

Further, chromatin accessibility was GRA28-dependently elevated around the promoter of *Il12p40* (**Figure 6E**), and the promoter of *Ccl22*, a known target chemokine for GRA28 (57) (**Figure 6F**). In contrast, accessibility was not elevated for genes encoding cytokines *Ccl24*, *Tnf* and *Il1a* (**Figure S6C**). For DC-associated transcription factors, analyses showed GRA28-dependent elevated chromatin accessibility around the promoters of *Zbtb46* (**Figure 6G**) and *Batf3* (**Figure S7C**), while accessibility changes were less evident for *Irf4* (**Figure 6H**) and *Nr4a3* (**Figure S7C**). Finally, ATAC-seq detected minor changes for *Cd86* (**Figure 6I**) and we observed a decreased expression of CD86 in PRUΔ*gra28*-infected BMDMs compared with wild-type-infected BMDMs, determined by flow cytometry (**Figure 6J)**. We conclude that GRA28 selectively elevates chromatin accessibility around the promoters of *Ccr7, Il12b*, *Ccl22* and genes encoding DC-associated transcription factors, thereby impacting their transcription in *T. gondii*-infected BMDMs.

### Impact of GRA15/24 double deletion on *T. gondii* dissemination in mice and on phenotypes of human macrophages/monocytes

We previously established a role for GRA28 in the migration of infected macrophages to secondary organs (21). Given the collective contributions of GRA15/24/28 to gene expression leading to pro-migratory activation of macrophages *in vitro*, we addressed the impact of GRA15/24 signaling *in vivo* on parasite dissemination. Equivalent numbers of pre-labelled BMDMs challenged with wild-type-or Δ*gra15*Δ*gra24* parasites were adoptively transferred i.p. to mice in competition assays (**Figure 7A**). Fourteen-18 h post-inoculation, organs were harvested and cells were characterized by flow cytometry (**Figure 7B** and **S8A**). Importantly, BMDMs challenged with *Δgra15Δgra24 T. gondii* migrated at a relative lower frequency to the omentum, mesenteric lymph nodes (MLNs) and spleen, compared with BMDMs challenged with wild-type *T. gondii* (**Figure 7C**), showing an implication of GRA15/24 in the migration of parasitized BMDMs in mice.

**Figure 7.**
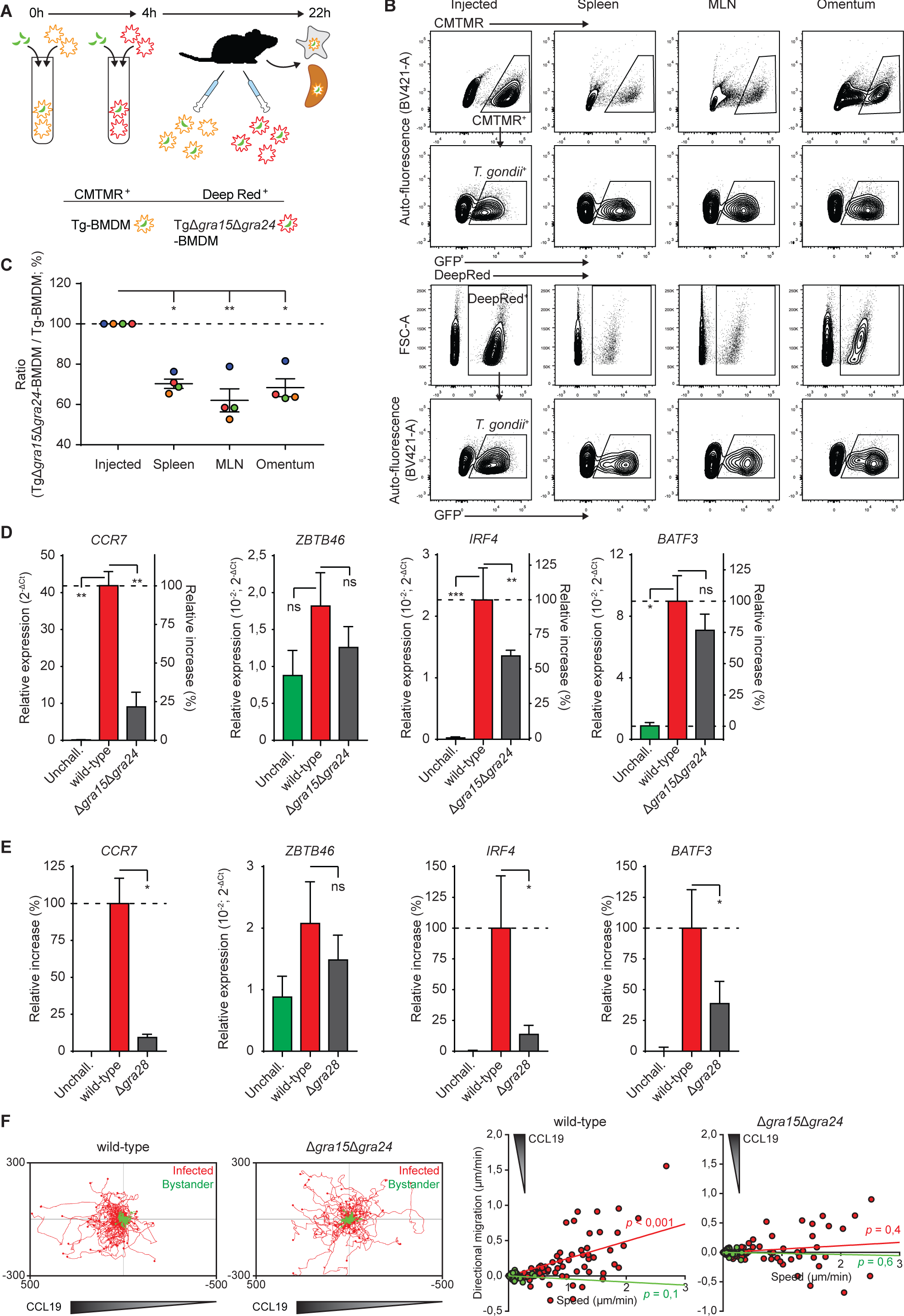
Impact of GRA15/24 on T. gondii dissemination in mice and on the phenotypes of human macrophages and monocytes. (**A**) Illustration of experimental setup and conditions for co-adoptive transfers of BMDMs challenged with *T. gondii* (PRU) wild-type (Tg-BMDM) or GRA15/24 double mutant (TgΔ*gra15*Δ*gra24*-BMDM) and pre-labeled with CMTMR or CellTracker Deep Red dyes, respectively. (**B**) Contour plots show typical gating strategy for flow cytometric detection of pre-labelled and *T. gondii* parasitized BMDMs (CMTMR^+^/Deep red^+^ and GFP^+^) as injected intraperitoneally and extracted from spleen, mesenteric lymph nodes (MLN) and omentum 18h post-inoculation, as detailed under Methods. Single (CD11c^+^) BMDMs were pre-gated as shown in **Figure S8A**. (**C**) Flow cytometric analysis of wild-type- or Δ*gra15*Δ*gra24*-challenged BMDMs in the spleen, MLNs and omentum 18h post-inoculation. Data is presented as the change in ratio between detected challenged Deep red^+^GFP^+^ (Δ*gra15*Δ*gra24*) cells and CMTMR^+^GFP^+^ (wild-type) cells related to the inoculated ratio (normalized to 100%). Mean ratio change ±SE and individual mice (n = 4) are displayed. (**D**) and (**E**) qPCR analyses of *Ccr7*, *Il12p40*, *Zbtb46* and *Irf4* cDNA from human monocyte-derived macrophages (mo-macs) challenged with *T. gondii* type II PRU (wild-type), (D) Δ*gra15*Δ*gra24* or (E) Δ*gra28* tachyzoites (18h, MOI 2). Displayed is relative expression (2^-ΔCt^) or the relative and increase in expression relative to untreated unchallenged (unchall., 0%) and wild-type (100%) challenged conditions (mean + SEM, n=4). (**F**) Motility plots depict the displacement of mo-macs challenged with wild-type and Δ*gra15*Δ*gra24* tachyzoites (14h MOI 1) in a CCL19 gradient as detailed in Methods (scale indicates µm; n=3). For each condition, directional migration (µm/min) towards the CCL19 source and speed (µm/min) of individual cells are displayed in graphs, with linear regression lines. Infected cells (GFP^+^, red) were analyzed. For each condition, p-values indicate the directional migration compared to hypothetical zero directionality (one-sample permutation test). Statistical comparisons were made with WLS and Dunnett’s post-hoc tests (C) or ANOVA and Dunnett’s post-hoc tests (D, E * p ≤ 0,05, ** p ≤ 0,01, ** p ≤ 0,001, ns p > 0,05).

Further, we extended our key findings in murine cells by challenging human peripheral blood monocytes or monocyte-derived macrophages (mo-macs) with two separate type II *T. gondii* lines (PRU or ME49-PTG). *T. gondii*-challenged cells expressed elevated levels of *CCR7*, *ZBTB46*, *IRF4* and *BATF3* compared with their unchallenged counterparts (**Figure 7D, Figure S8B**). Similar to murine BMDMs, challenge with GRA15/24-deficient tachyzoites yielded decreased expressions (**Figure 7D**), in line with effects of GRA28-deficiency (**Figure 7E**). Finally, we confirmed abolished chemotaxis in a CCL19 gradient by mo-macs challenged with GRA15/24-deficient parasites (**Figure 7F**), with maintained hypermotility (**Figure S8C)**. Jointly, we conclude that GRA15/24/28 potentiate the migration of infected macrophages to secondary organs in mice and induce pro-migratory transcriptional activation and CCR7-driven chemotaxis in human macrophages and monocytes.

## Discussion

Here, we addressed how *T. gondii* orchestrates the migratory activation of macrophages that promotes systemic parasite dissemination. We demonstrate that a set of secreted GRA proteins regulates the chemotactic and pro-inflammatory activation of parasitized macrophages. First, we identify novel functions for GRA15 and GRA24 in promoting CCR7-mediated chemotactic responses. Second, we show that GRA15 and GRA24 cooperate by acting on NF-κB and p38 MAPK signaling pathways, respectively, with minor contributions by GRA16 and GRA18 and counteracting effects by TEEGR to the migratory and pro-inflammatory responses of parasitized macrophages. Third, we report that GRA28 elevates chromatin accessibility at the *Ccr7* locus and other loci associated with pro-migratory activation of macrophages. Finally, we show that GRA15/24 impact systemic transport of *T. gondii* in mice, similar to GRA28 (21). We propose a model for how the concerted action of GRA effectors activates parasitized macrophages (**Figure 8**).

**Figure 8.**
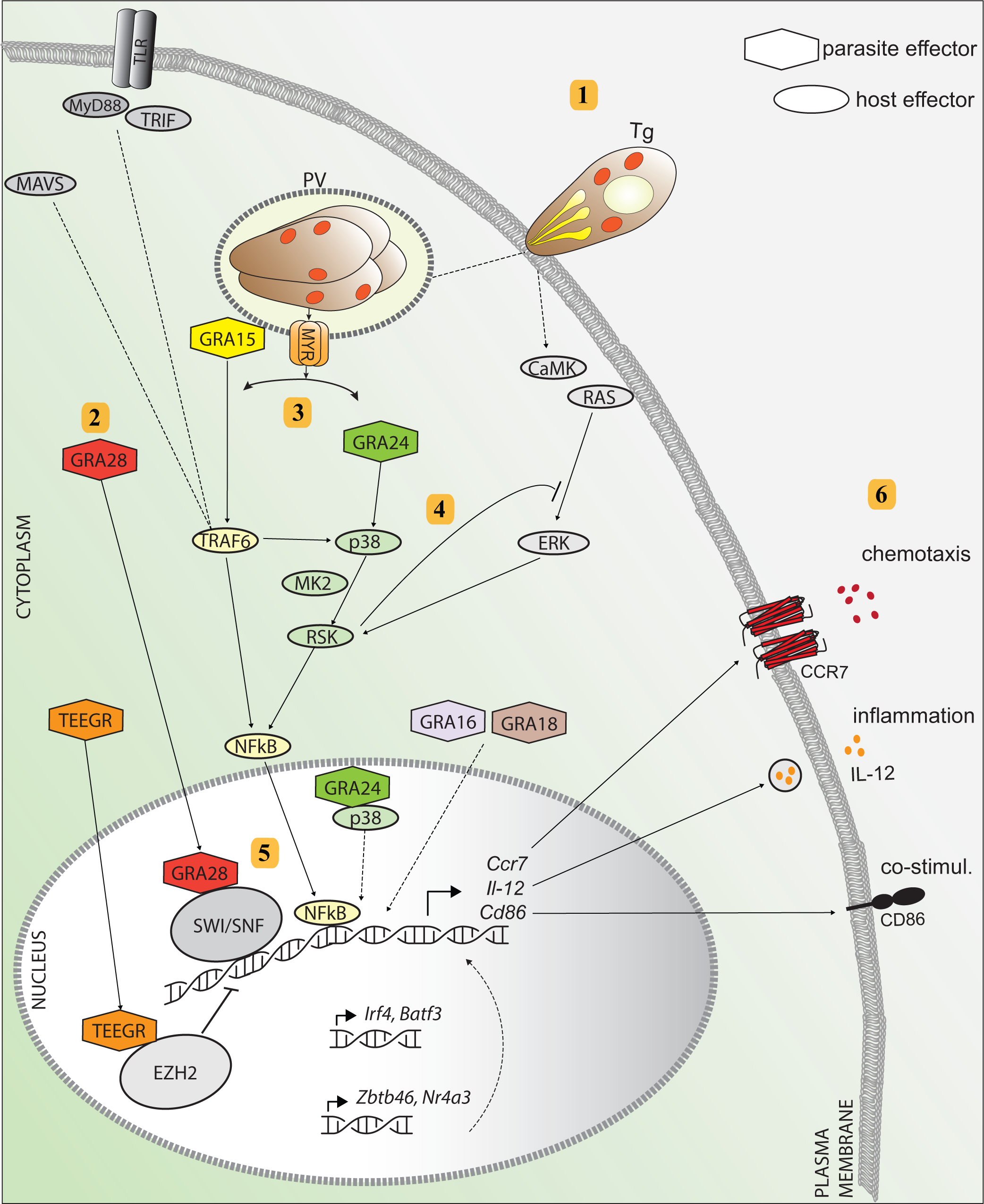
Hypothetical model for the migratory activation of parasitized macrophages by Toxoplasma. **1.** *T. gondii* actively invades the macrophage and forms a parasitophorous vacuole (PV) where it replicates and secretes effector proteins from secretory organelles (dense granules) into the host cell cytosol via the MYR1 translocon. **2.** The effector GRA28 is secreted into the host cell cytosol MYR1-dependently and locates to the nucleus where it complexes with chromatin remodelers (SWI/SNF) to open up chromatin. **3.** GRA15 is secreted MYR1-independently and interacts with TRAF6 to activate NF-κB, which locates to the nucleus. In parasitized macrophages, NF-κB activation occurs independently of TLR/MyD88 /TRIF/MAVS signaling. **4.** MYR1-dependent secretion of GRA24 activates p38 signaling and potentiates NF-κB activation via RSK and MK-2. The GRA24/p38 complex can also travel to the nucleus. **5.** GRA28-mediated increased chromatin accessibility facilitate GRA15/24/p38-driven transcription via NF-κB at the *Ccr7, Il12 and Cd86* gene loci. GRA16 and GRA18 contribute to these effects by unknown mechanisms. The effector TEEGR counteracts these effects, presumably through interaction with the transcriptional repressor EZH2. The elevated transcription of DC-related transcription factors (*Zbtb46*, *Irf4*, *Nr4a3*, *Batf3)* also drives expression of *Ccr7, Il12 and Cd86*. **6.** Altered signaling results in elevated expression of CCR7 with chemotactic responses, pro-inflammatory IL-12 and CD86 expression.

We report a role for GRA15 in the migratory activation and CCR7-driven chemotaxis of parasitized macrophages via its activation of NF-κB signaling (30). The findings have a direct bearing on the *Trojan horse* mechanism for *T. gondii* dissemination also because GRA15 impacts ICAM-1 expression (58) and adhesion molecules regulate the motility and migration modes of macrophages in tissues (59). In line with this, we recently reported that DCs infected with GRA15-deficient *T. gondii* exhibit reduced transmigration across endothelium (58). However, GRA15-deficiency reduced, but did not strictly abolish, transcriptional activations and the migratory phenotypes of macrophages and DCs. Moreover, the migratory phenotypes are present in the type I RH line, which lacks a functional GRA15 (60). Thus, while GRA15 likely contributes to the reported enhanced transmigration frequencies of type II-infected DCs over type I-infected DCs (12), additional effectors are in play in a strain/genotype dependent manner.

We demonstrate a role for GRA24 in the migratory activation of infected macrophages, mediated via p38 MAPK signaling. In type II *T. gondii*-infected macrophages, this response was prominent and co-operated with GRA15 in eliciting chemotactic and pro-inflammatory activation. Consistent with this, GRA15 and GRA24 were recently reported to synergistically promote pro-inflammatory cytokine expression in type II strains (61). Interesting findings in this context are the impact of GRA15 on p38 phosphorylation and the p38-linked activation of RSK, jointly advocating for cross-regulation between NF-κB and p38 MAPK signaling. Consequently, a GRA15/24 double mutant dramatically reduced the migratory activation of macrophages *in vitro* and *in vivo*, corroborating the importance of these two parasite effectors. Similarly, combined inhibition of NF-κB and p38 MAPK signaling mirrored these effects. Yet, in neither case was the phenotype totally abrogated, indicating the implication of additional effectors or signaling. Interestingly, the dense granule protein TEEGR acted as a down-modulator of *Ccr7, Il12p40* and other responses, presumably related to its negative regulation of NF-κB (52). In contrast, GRA16 and GRA18 positively contributed to the phenotype, however to a lesser extent than GRA15 or GRA24. Along these lines, in type I (RH) *T. gondii*-infected macrophages, we found that the responses, specifically CCR7 chemotaxis, were less dependent on GRA24 (21). One explanation might be that, because the type I RH strain lacks a functional GRA15 protein that activates NF-κB responses (30), NF-κB activation is mediated by other GRA or non-GRA effectors and that the balance between NF-κB activating and counteracting effectors is shifted (52, 62, 63). Finally, the comparable induction of *Ccr7, Il12p40, Zbtb46* and *Irf4* expression observed in both wild-type and Myd88^-/-^ Ticam^-/-^ Mavs^-/-^ macrophages (49) indicates that, upon *T. gondii* infection, activation of NF-κB signaling predominantly occurs in a manner that is independent of the MyD88/TRIF/IPS-1-mediated pathways for NF-κB activation. Instead, the mounting data indicates that the activation of NF-κB for this phenotype occurs via TRAFs (30, 47).

We demonstrate that GRA28 selectively elevates chromatin accessibility at the *Ccr7* gene locus. This effect, in conjunction with GRA28’s interaction with the chromatin remodeler SWI/SNF (21), establish a central role for GRA28 in the migratory activation of macrophages by both type I and II *T. gondii* strains. We propose a mechanistic framework in which GRA28 interacts with chromatin remodelers to fine-tune the transcriptional regulation directed by GRA15—NF-κB and GRA24—p38 MAPK, targeting the expression of *Ccr7, Il12p40* and *Cd86* (**Figure 8**). Further corroborating this model, the *Ccr7* promotor contains 4 putative NF-κB binding sites and a single putative AP-1/c-Fos binding site (64). Pharmacological inhibitor data indicate that NF-κB, but not AP-1 or PU.1, is the major transcription factor in play. Jointly, the findings establish a central role for the GRA15/24/28 triad in the migratory activation of macrophages, supporting previous observations that genotype-specific parasite effectors influence *T. gondii* dissemination by parasitized leukocytes (12). GRA28 selectively enhanced chromatin accessibility at the *Ccr7, Il12p40* and *Ccl22* gene loci, without a detectable enhancement at gene loci of other important cytokines, chemokines or chemokine receptors. Thus, the molecular determinants guiding the site-specificity of GRA28 or GRA28/SWI/SNF complexes at these loci warrant further investigation.

Why does *T. gondii* express and secrete multiple effectors that co-operate in the migratory activation of phagocytes? *T. gondii* infects and replicates within different types of immune cells in a broad array of vertebrate species. Seeking latency in warm-blooded vertebrates to assure later transmission to the feline definitive host necessitates dissemination to various organs, especially the brain, which must be executed across a range of host species and immune cell types. Moreover, different anatomical sites and temporal stages of infection might necessitate distinct migratory behaviors such as reverse transmigration, afferent migration, and motility in tissues. Thus, polymorphisms and functional redundancy in effector proteins may enable *T. gondii* to adapt to both inter-host and intra-host diversity. Our current understanding points towards a nuanced activation of different phagocytic cell types. Specifically, while DCs are readily chemotactically activated (22, 23), macrophages exhibit a relative resilience against such activation, as demonstrated by their lack of CCR7 upregulation or chemotaxis upon PRR stimulation (21, 65). GRA28 appears instrumental in overcoming this resilience by mediating chromatin accessibility to the *Ccr7* locus thus enabling NF-κB-induced activation where it would otherwise not occur.

GRA15/24/28 target complementary host pathways eventually leading to CCR7 expression and migratory activation. It is reasonable to speculate that such system redundancy ensures the robustness of leukocyte hypermigration, a hypothesis supported by existing literature (66). Further, the diversity in effector molecules could potentially aid in evading host immune recognition, thereby thwarting efforts at neutralization. On the other hand, hypermigration of phagocytes might also increase contact between immune cells, for example, with the T cell compartment (67), potentially leading to increased immune control, albeit, without hindering chronic infection. Lastly, the observed counteractive interactions among various GRA and ROP proteins, such as GRA15 vs TEEGR for NF-κB activation or ROP16 vs GRA28 for CCR7 regulation, hint at an additional layer of complexity, possibly functioning as a fine-tuning mechanism or as on-off switches. Therefore, it can be postulated that effector diversification serves as a flexible strategy for the parasite, allowing for both stringent regulation and evolutionary adaptability across multiple hosts.

Microbial pathogens have developed elaborated strategies to thrive inside phagocytes and colonize their hosts (68-70). For example, recent findings show that *Mycobacterium tuberculosis* manipulates alveolar macrophage trafficking for rapid localization to the lung interstitium (71). Further, *Leishmania* and *Salmonella typhimurium* inhibit macrophage and DC motility (72, 73) while coccidian parasites elevate the migration of phagocytes (13). However, a detailed molecular understanding of how pathogens orchestrate the hijacking of complex cellular processes, such as host cell migration, is yet to come. The present findings provide a molecular framework delineating how *T. gondii* orchestrates the migratory activation of phagocytes, primarily via the concerted action of GRA15/24/28 targeting CCR7-mediated chemotaxis. Yet, hypermotility is maintained in GRA15/24/28 mutants (21, 37). Indeed, in the hypermigratory responses of phagocytes (17), separate but partly overlapping signaling regulate GABA/voltage-dependent calcium channel (VDCC)-driven hypermotility and CCR7-driven chemotactic activation of parasitized phagocytes. Specifically, the rhoptry protein TgWIP modulates the host cell actin dynamics presumably via WAVE/arp2/3 complex (20), while ROP17 presumably acts on RhoGTPases (19) and via the MYR1 complex (21, 74). Also, Tg14-3-3-related sequestration of host 14-3-3 to the PV presumably impacts MAPK signaling (16, 18). However, the specific parasite-derived effector(s) activating motogenic GABAergic and VDCC-mediated signaling remain to be identified (13). In summary, we propose that the joint activities of these effectors culminate in pro-migratory signaling within the infected phagocyte. Such coordinated action may not only facilitate the parasite’s dissemination within multiple hosts but could also confer advantages in evading host immune responses. The implications of this pro-migratory signaling state and its ultimate consequences for host-pathogen interactions merit further in-depth investigation.

## Acknowledgements

We are grateful to Dr Nelson Gekara, Stockholm University and University of Freiburg, for providing Myd88^-/-^ Ticam^-/-^ Mavs^-/-^ mutant material. This paper benefited from publicly available data generated by the Immunological Genome Project Consortium (ImmGen).

The studies were funded by the Swedish Research Council, grants 2018-02411 (A.B.), 2022-00520 (A.B.), 2020-03818 (D.M.O), NIH grant R01 AI166715 (J.P.J.S.), Carl Tryggers Foudation CTS 21:1158 (D.M.O), Åhlen Foudation grant 223020 (A.B.) and the Sven and Lily Lawski Foundation (M.E.R./A.B.).

The authors declare that they have no conflicts of interests.

## Materials and methods

Cell lines, mouse strains and parasite strains are described in **Table S1**. Antibodies, chemicals and kits are detailed in **Table S2**.

### Ethical considerations

The Regional Animal Research Ethical Board, Stockholm, Sweden, approved experimental procedures and protocols involving extraction of cells from mice (permit numbers 14458/2019 and 16334-2022), following proceedings described in EU legislation (Council Directive 2010/63/EU). The Regional Ethics Committee, Stockholm, Sweden, approved protocols involving human cells (application number 2006/116-31). All donors received written and oral information upon donation of blood at the Karolinska University Hospital Blood Center. Written consent was obtained for utilization of white blood cells for research purposes.

### Mouse cell culture

Cells from bone marrow of 6-10-week-old male or female wild-type or Myd88^-/-^ Ticam^-/-^ Mavs^-/-^ (49) C57BL/6 mice (see **Table S1**) were cultivated in RPMI 1640 (VWR) with 10% fetal bovine serum (FBS; HyClone), gentamicin (20 μg/mL; Sigma-Aldrich) and glutamine (2 mM), referred to as complete medium, and supplemented with 20 ng/mL recombinant mouse GM-CSF. Strongly adherent cells were harvested on day 6-8 as bone marrow-derived macrophages (BMDMs). For peritoneal macrophages (PEMs) C57BL/6 mice were euthanized and peritoneal lavage (10 ml PBS) was collected from the peritoneal cavity. After overnight culture in complete medium, loosely and non-adherent cells were removed by repeated washing and the adherent cells were used as PEM in experiments for RNA isolation.

### Human cell culture

Human CD14^+^ monocytes were isolated from peripheral blood mononuclear cells (PBMC) after density gradient centrifugation on Lymphoprep with CD14 MicroBeads from buffy coats obtained from healthy donors at the Karolinska University Hospital Blood Center and cultured in complete medium. Monocyte-derived macrophages (mo-macs) were generated from CD14^+^ monocytes through culture for 3-5 days in complete medium supplemented with 20 ng/mL human recombinant GM-CSF.

Human foreskin fibroblasts HFF-1 were cultured in Dulbecco‘s modified Eagle’s medium, high glucose, (DMEM; VWR) with 10% fetal bovine serum (FBS; HyClone), gentamicin (20 μg/ml; Sigma-Aldrich) and glutamine (2 mM) and HEPES (0.01 M), referred to as DMEM.

### Parasite culture

*T. gondii* tachyzoites were maintained by serial 2-day passages in human foreskin fibroblast HFF-1 monolayers. Freshly egressed tachyzoites were used for all infections. The different strains used are listed in the **Table S1**. All cell cultures used were periodically tested for mycoplasma and found to be negative.

### Infection challenges

Carry-over from routine *T. gondii* culture to experiments was minimized by repeated washing of the freshly egressed tachyzoites before challenge with live tachyzoites. For all qPCR, ATAC-seq and western blot experiments, BMDMs, human monocytes, mo-macs and/or PEMs were challenged with freshly egressed *T. gondii* tachyzoites with indicated strains/lines at MOI 2 for 18h, unless differently stated. LPS was used at a final concentration of 10 ng/mL. For flow cytometry assays, BMDMs were challenged for 18h with GFP-expressing or CFSE-labeled *T. gondii* tachyzoites with the indicated strains (MOI 1). For chemotaxis experiments, BMDMs were challenged for 14h with GFP-expressing or CFSE-labeled *T. gondii* tachyzoites at MOI 1 before seeding in the chemotaxis chamber.

### Inhibitors

When indicated, cells were treated with inhibitors Trametinib (1 µM), BIRB 796 (10 µM), JNK-IN-8 (3 µM), BRD7389 (5 µM), MK2-IN-1 (5 µM), TPCA-1 (3 µM), SR 11302 (20 µM), T-5224 (2 µM), DEL-22379 (5 µM), DB2313 (1 µM), BIX02189 (0,5 or 1 µM) and/or JSH-23 (25 µM) vehicle (DMSO) initiated 1h prior to challenge.

### Quantitative polymerase chain reaction (qPCR)

BMDMs, PEMs and human monocytes and mo-macs were cultured with complete medium or challenged with freshly egressed *T. gondii* tachyzoites of the indicated strains and lysed in TRI Reagent (Sigma-Aldrich) or Lysis buffer (Jena Bioscience). Total RNA was extracted according to the manufacturers’ protocol using the Direct-zol RNA Miniprep (Zymo Research) or Total RNA Purification (Jena Bioscience) kits and reverse transcribed with Maxima H Minus Reverse Transcriptase (Thermo Fisher). Real time qPCR was performed with SYBR® green PCR master mix (KAPA biosystems) or HotStart™ 2X SYBR Green qPCR Master Mix (APExBIO Technology), specific forward and reverse primers at target-dependent concentrations (100-200 nM) and cDNA (10-30 ng) in a QuantStudio 5 System (Thermo Fisher) with ROX as a passive reference. qPCR results were analyzed using the ΔCq method relative to Importin-8 and TATA-binding protein (TBP) as housekeeping genes and displayed as such or normalized to unchallenged (set to 0%) and wild-type *T. gondii*-challenged (set to 100%). Primers are listed in **Table S3**.

### Assay for transposase-accessible chromatin with sequencing (ATAC-seq)

To measure chromatin accessibility, we performed ATAC-seq using ATAC-Seq Kit according to manufacturer’s instructions. After challenging macrophages with *T. gondii*, we centrifuged 100,000 cells at 500 × *g* at 4°C for 5 min, removed the supernatant, and washed the cells once with 100 µl ice-cold 1 × PBS without disturbing the cell pellet. We next performed additional centrifugation at 500 × *g* at 4°C for 5 min. Afterwards, we removed 1 × PBS and resuspended the cells in 100 µl ATAC lysis buffer which is immediately followed by centrifugation at 500 × *g* at 4°C for 10 min. After removing the supernatant, we incubated the nuclei in 50 µl Tagmentation Master Mix (25 µl 2 X Tagmentation buffer, 12 µl water, 10 µl Assembled Transposomes, 2 µl 10 X PBS, 0.5 µl 1% Digitonin, 0.5 µl 10% Tween 20) at 37°C for 30 min using PCR machine with heated lid. Following the incubation, we added 250 µl DNA binding buffer and 5 µl Sodium acetate and performed column purification by centrifugation at 17,000 x *g* for 1 min. The column was washed once with 750 µl wash buffer and centrifuged at 17,000 x *g* for 1 min. Finally, we eluted tagmented DNA from the column using 35 µl DNA Purification Elution Buffer by centrifugation at 17,000 x *g* for 1 min.

### PCR amplification of tagmented DNA

33.5 µl previously isolated tagmented DNA is mixed with 10 µl 5 × Q5 Reaction buffer, 2.5 µl 25 µM i7 Indexed primer, 2.5 µl 25 µM i5 Indexed primer, 1 µl 10mM dNTPs and 0.5µl 2 U/µl Q5 Polymerase. The reaction mixture was then incubated on a thermocycler with the following PCR parameters: 72°C for 5 min, 98°C for 30 sec, 10 cycles of 98°C 10 sec, 63°C 30 sec, 72°C one min. ATAC-seq libraries were sequenced as 79 + 79 nt paired end reads using NextSeq 550 (Illumina).

### Analysis of ATAC-seq data

ATAC-seq dataset was aligned via Bowtie 2.2.5 (75) with parameters “--very-sensitive --no-unal --no-mixed --no-discordant -I 10 -X 700”. The alignment results were converted to bigWig signal tracks using BEDtools 2.27.1 and SAMtools (76) and normalized to uniquely aligned reads. The signals (log10 RPM with pseudo count 0.01) at TSS+-500bp were also calculated for each sample and used for generating Pearson correlation co-efficiency. MACS2 (77) was then applied to call peaks with default parameters. Peaks present in both independent biological replicates were regarded as representative peaks for each condition and used for generating the Venn diagram.

### Flow cytometry

BMDMs were challenged as indicated and stained with anti-CD11c, CD11b, MHCII I-A/I-E, CD86, or isotype control antibodies and live/dead far red stain. Staining was performed on fixed (2% PFA) or live cells, blocked with anti-CD16/CD32 antibody, in FACS buffer (PBS/0,5% FBS/1mM EDTA). Flow cytometry was performed on a BD LSRFortessa flow cytometer (BD Biosciences) and analyzed with FlowJo X (FlowJo LLC).

### Western blot

For Western blotting cells were challenged as indicated, harvested, washed with PBS and then lysed directly in Laemmli buffer for whole cell lysates. Proteins were separated using 10% SDS-PAGE, blotted onto a PVDF membrane and blocked (10% BSA in TBS/0,5% Tween-20) followed by incubation with primary and secondary antibodies: anti-TATA-binding protein, anti-p-IκBα (Ser32/36), anti-p-RSK (Ser380/386), anti-p-p38 (Thr180/Tyr182) anti-mouse, anti-rabbit or anti-rat IgG-HRP in 5% BSA/0,5% Tween-20 in TBS. Proteins were revealed by mean of enhanced chemiluminescence (GE Healthcare) in a BioRad ChemiDoc XRS+. Densitometry analysis was performed using ImageJ (NIH, MD, USA). For display, the contrast of images was enhanced with the ImageJ ‘enhance contrast’ feature without pixel saturation.

### Chemotaxis

BMDMs or human macrophages were challenged as indicated, washed, resuspended in CM with 1 mg/mL bovine collagen type I (Sigma) and seeded into uncoated ibiTreat μ-slide chemotaxis chambers (Ibidi, Martinsried, Germany). Collagen was allowed to polymerize for 30 min and media, inhibitors and 1,25 µg/mL murine (BMDMs) or human recombinant CCL19 (human macrophages) were added as indicated conform the manufacturer’s instructions (application note 23). Cells were then imaged every 5 min for 8h (Zeiss Observer Z.1). Motility tracks for ≥35 cells per condition were analyzed using ImageJ software for each experiment.

### Immunofluorescence microscopy

BMDMs were seeded on gelatin (1%)-coated glass coverslips, challenged freshly egressed *T. gondii* tachyzoites (PRU A7; MOI 2) for 18h or left unchallenged for 6h (p-RSK) and fixed with 2% PFA. Cells were then permeabilized with 0,1% Triton X-100 in PBS and stained with phalloidin Alexa Fluor 594 (Thermo Scientific) or primary anti-p-RSK (Ser380/386) and anti-rabbit IgG Alexa Fluor 594 (Thermo Scientific) secondary antibodies and DAPI. Images were acquired on a Leica DMi8 with 63x objective.

### Adoptive transfers

Adoptive transfers were performed and analyzed as previously described (21). Briefly, BMDMs were stained with CellTracker CMTMR (2 μM) or Deep Red (1 μM) dyes (2,5x10^6^ cells each), washed and challenged with indicated freshly egressed *T. gondii* GFP-expressing or CFSE-stained type II tachyzoites (PRU; MOI 1,5) in complete medium for 4h. Cells were then washed and injected i.p. into C57BL/6 mice. Mice were sacrificed 18h post-injection to collect spleens, mesenteric lymph nodes and omenta. The organs were triturated, filtered through 40 µm cell strainer and fixed (4% PFA). Cells from the spleen were blocked with anti-CD16/32 antibody in FACS buffer (PBS/0,5% FBS/1mM EDTA) and stained with CD11c antibody. Samples were then analyzed by flow cytometry on a BD LSRFortessa flow cytometer (BD Biosciences) and with FlowJo X (FlowJo LLC).

### Handling of publicly available datasets

ChIP- and ATAC-seq data available from NCBI GEO series GSE100738 (ATAC-seq), GSE57563 (H3K4me1) and GSE64767 (H3K4me3), which are described elsewhere, was visualized in the USCS genome browser (56, 78). RNA-seq data is taken from the ImmGen Ultra-low-input RNA-seq dataset, NCBI GEO superseries GSE127267.

### Statistical analyses

All experiments were replicated to allow for statistical comparisons as stated for each experiment in the figure legends. The number of n denotes the number of biological replicates (adoptive transfer experiments: individual mice) or independent experiments (all other experiments). Statistical analyses were performed with R, RStudio and packages afex (repeated-measures ANOVA), nlme (Weighted Least Squares), emmeans (Dunnett’s post-hoc), DAAG and rcompanion (permutation tests). Hypothesis tests and inferential statistics used are indicated in the figure legends and were chosen based on experimental design, the hypothesis to be tested and data distribution and which statistics were to be presented. In cases of heteroscedascity due to normalization, multiple comparisons were done after Weighted Least Squares (WLS) regression, otherwise ANOVA was run. Dose-dependent inhibition was tested with Spearman rank correlation. Chemotaxis analyses were done with linear regression for visualization of regression lines and, due to non-normal distribution with one-sample permutation tests for hypothesis testing. Reported p-values from multiple comparisons were corrected with the Holm-Bonferroni method. Statistical significance is defined as p < 0,05.

## Supplementary figures

**Figure S1. Phenotypical and transcriptional responses of BMDMs to *T. gondii*-challenge**

(**A**) Frequency of infected (GFP^+^) cells among BMDMs challenged with freshly egressed GFP-expressing *T. gondii* type II wild-type (WT) or GRA15-deficient (Δ*gra15*) tachyzoites for 18 h (PRU, MOI 2; n=3).

(**B**) Motility of BMDMs challenged with freshly egressed *T. gondii* type II GRA15-deficient (PRUΔ*gra15*) tachyzoites over 14-16 h in a collagen matrix with a CCL19 gradient as detailed in Methods (scale indicates µm). Dots indicate mean speed of individual cells and lines and error bars are mean ± SEM.

(**C**) Gating strategy for flow cytometric analysis of infected (GFP^+^) and bystander (GFP^-^) BMDMs challenged with freshly egressed GFP-expressing *T. gondii* type II tachyzoites.

(**D**) and (**E**) Flow cytometric analysis of anti-CD40 and CD80 (D) or MHCII (E) staining on BMDMs challenged for 18 h with freshly egressed GFP-expressing *T. gondii* type II wild-type (WT) and GRA15-deficient (Δ*gra15*) tachyzoites (PRU; MOI 1) or left unchallenged. Infected (GFP^+^) and bystander cells (GFP^-^) were analyzed. Bar graph displays the MFI of infected and unchallenged conditions (mean + SEM; n=5).

(**F**) qPCR analysis of *Egr1* cDNA from BMDMs challenged for 18 h with freshly egressed *T. gondii* type II wild-type and GRA15-deficient (Δ*gra15*) tachyzoites (PRU; MOI 2) or left unchallenged (unchall.). Displayed are relative expression (2^-ΔCt^) and the increase in expression relative to unchallenged (0%) and wild-type (100%) conditions (mean + SEM; n=5).

(**G**) qPCR analysis of *Ccr7, Il12p40, Zbtb46* and *Irf4* cDNA from BMDMs challenged for 6 h with freshly egressed *T. gondii* type II wild-type tachyzoites (PRU; MOI 2) in the presence (IKKi) or absence of TPCA-1 (-) or left unchallenged (unchall.). Displayed are relative expression (2^-ΔCt^) and the increase in expression relative to unchallenged (0%) and wild-type (100%) conditions (mean + SEM; n=4).

Statistical comparisons were made with paired t-test (A), pairwise permutation test (B) or ANOVA and Dunnett’s post-hoc tests (D-H; * p ≤ 0,05, ** p ≤ 0,01, *** p ≤ 0,001, ns p > 0,05).

**Figure S2. Roles of MAP kinases, AP-1 and PU.1 in the transcriptional activation of T. gondii challenged macrophages**

(**A**) qPCR analysis of *Ccr7, Il12p40, Zbtb46* and *Irf4* cDNA from BMDMs challenged for 18 h with freshly egressed *T. gondii* type II GRA24-deficient tachyzoites (PRUΔ*gra24*; MOI 2) in the absence (-) or presence of MEK1/2, p38 MAPK or JNK inhibitors or left unchallenged (unchall.). Displayed is the increase in expression relative to unchallenged (0%) and *T. gondii* type II wild-type-challenged (100%) conditions (mean + SEM; n=4).

(**B**) qPCR analyses of *Ccr7, Il12p40, Zbtb46* and *Irf4* cDNA from BMDMs challenged for 18 h with *T. gondii* type II tachyzoites (PRU, MOI 2) in the absence (-) or presence of ERK1/2 dimerization inhibitor DEL-22379, CaMKK inhibitor STO-609 (CaMKKi) or left unchallenged (unchall.). Displayed is the increase in expression relative to unchallenged (0%) and wild-type-challenged (100%) conditions (mean + SEM; n=3 (DEL-22379) or 4 (STO-609)).

(**C**) Motility plots depict the displacement of BMDMs challenged with freshly egressed *T. gondii* type I tachyzoites (RH1-1; MOI 1) in the presence or absence (vehicle) of p38 MAPK (p38i) or MEK1/2 (MEKi) over 14 h in a collagen matrix with a CCL19 gradient as detailed in Methods (scale indicates µm). Infected cells (GFP^+^) were analyzed.

(**D**) Representative micrograph shows unchallenged BMDMs stained for p-RSK (S380/386, red) and nuclei (DAPI, blue). Scale bar = 10 µm.

(**E**) Western blot analysis of p-RSK (S380/386) expression in cytoplasm- and nucleus-enriched fractions of BMDMs challenged for 5 h with wild-type and GRA24-deficient (Δ*gra24*) *T. gondii* type II tachyzoites (PRU, MOI 3).

Statistical comparisons were made with ANOVA and Dunnett’s post-hoc tests (A) or paired t-test (B, C; * p ≤ 0,05, ** p ≤ 0,01, ns p > 0,05).

**Figure S3. Transcriptional impacts of AP-1 and PU.1 inhibition on DMDMs**

(**A**) and (**B**) qPCR analyses of *Ccr7, Il12p40, Zbtb46* and *Irf4* cDNA from BMDMs challenged for 18 h with *T. gondii* type II tachyzoites (PRU, MOI 2) in the absence (-) or presence of (A) AP-1 inhibitors SR 11302 (SR) and T-5224 (T5) or (B) PU.1 inhibitor DB2313. Displayed is the increase in expression relative to unchallenged (0%) and wild-type-challenged (100%) conditions (mean + SEM; n=3 (B) or 4(A)). Statistical comparisons were made with ANOVA and Dunnett’s post-hoc tests (A) or paired t-test (B; * p ≤ 0,05, ns p > 0,05).

**Figure S4. Responses of Myd88^-/-^ Ticam^-/-^ Mavs^-/-^ macrophages to LPS**

qPCR analysis of *Il12p35* and *Il12p40* cDNA from wild-type (WT) or Myd88^-/-^ Ticam^-/-^ Mavs^-/-^ (TKO) BMDMs challenged for 18 h with LPS (10 ng/mL) or left unchallenged (unchall.). Displayed is relative expression (2^-ΔCt^) (mean + SEM; n=2).

**Figure S5. Impact of GSK3β inhibition on the transcriptional activation of macrophages**

qPCR analyses of *Ccr7*, *Il12p40*, *Zbtb46* and *Irf4* cDNA from BMDMs challenged with *T. gondii* wild-type or GRA18-deficient (Δ*gra18*) tachyzoites (PRU), with or without AR-A014418 (GSK3i), as in (A). Bar graphs display the increase in expression relative to untreated unchallenged (unchall., 0%) and GRA18-deficient or wild-type (100%) challenged conditions (mean + SEM). Statistical comparisons were made with Student’s t-tests for paired samples (n=5, * p ≤ 0,05, ** p ≤ 0,01, ns p > 0,05).

**Figure S6. Gene expression and chromatin state in DCs and macrophages**

(**A**) Genome tracks show peak signal intensity (y-axis) for open or closed chromatin regions and indicated histone marks at selected genes from publicly available ATAC-seq and ChIP-seq data. Bar graphs show the corresponding mRNA expression of these genes from publicly available RNA-seq data. ATAC-seq tracks are of splenic CD8+ and CD4+ DC (cDC1/cDC2), peritoneal (MΦ PC) and alveolar macrophages (MΦ Alv). H3K4me1 (K4me1) and H3K4me3 (K4me3), indicative of active promoters and enhancers (79), ChIP-seq tracks are of *in vitro*-derived Flt3L-DC (cDC). See Materials and Methods for sources and details.

(**B**) Heatmap reports Pearson correlation co-efficient based on the chromatin accessibility measured by ATAC-seq between biological replicates of each condition: BMDMs challenged for 18h with *T. gondii* wild-type or GRA28-deficient (Δ*gra28*) tachyzoites (PRU) or left unchallenged.

**Figure S7. Chromatin accessibility and gene expression in DCs and macrophages**

(**A**) Genome tracks show peak signal intensity (y-axis) for open or closed chromatin regions of the *Ccr2*, *Ccr5* and *Cx3cr1* genes. For BMDMs, ATAC-seq signal from 2 separate biological replicates per condition. Upper tracks show peak signal from dendritic cells (cDC1) and peritoneal cavity macrophages (PC MΦ) extracted from Immgen publicly available dataset. Indicated is a region of interest (red outline) near the transcription start site (TSS).

(**B**) Visualization of open or closed chromatin regions of *Ccl24*, *Tnf* and *Il1a* genes as in (A).

(**C**) Visualization of open or closed chromatin regions of *Batf3* and *Nr4a3* genes as in (A) and mRNA expression from the publicly available Immgen RNA-seq dataset and qPCR presented in this paper. Statistical comparisons were made with ANOVA and Dunnett’s post-hoc tests (n=4-5, ** p ≤ 0,01).

**Figure S8. Characterizations of human monocytes and monocyte-derived macrophages**

(**A**) Gating strategy for flow cytometric detection in organs of intraperitoneally injected BMDMs (mesenteric lymph nodes, omentum) or CD11c^+^ BMDMs (spleen), based on gating of injected BMDMs as displayed. The histogram shows CD11c^+^ staining of cells extracted from spleen (blue) and injected BMDMs for reference (grey). The following gating steps are depicted in figure 7B.

(**B**) qPCR analyses of *Ccr7*, *Il12p40*, *Zbtb46* and *Irf4* cDNA from human monocytes challenged with *T. gondii* type II line ME49-PTG (18h, MOI 2). Displayed is relative expression (2^-ΔCt^) or the increase in expression relative to untreated unchallenged (unchall., 0%) and wild-type (100%) challenged conditions (mean + SEM, n=4).

(**C**) Motility of mo-macs challenged with PRU wild-type and Δ*gra15*Δ*gra24* tachyzoites (14h MOI 1) in a CCL19 gradient as detailed in Methods (scale indicates µm). Dots indicate mean speed of individual cells and lines and error bars are mean ± SEM. Statistical comparisons were made with pairwise permutation test (*** p ≤ 0,001).

